# Intrahost dynamics, together with genetic and phenotypic effects predict the success of viral mutations

**DOI:** 10.1101/2024.10.18.619070

**Authors:** Cedric CS Tan, Marina Escalera-Zamudio, Alexei Yavlinsky, Lucy van Dorp, Francois Balloux

## Abstract

Predicting the fitness of mutations in the evolution of pathogens is a long-standing and important, yet largely unsolved problem. In this study, we used SARS-CoV-2 as a model system to explore whether the intrahost diversity of viral infections could provide clues on the relative fitness of single amino acid variants (SAVs). To do so, we analysed ∼15 million complete genomes and nearly ∼8000 sequencing libraries generated from SARS-CoV-2 infections, which were collected at various timepoints during the COVID-19 pandemic. Across timepoints, we found that many of the SAVs that went on to reach high frequency could be detected in the intrahost diversity of samples collected at a median of 3-22 months prior. Additionally, we found that genetic linkage patterns observed at the interhost level can also be observed in the intrahost diversity of infections. Application of machine learning models allowed us to learn highly generalisable intrahost, physiochemical and phenotypic patterns to forecast the future fitness of intrahost SAVs (*r*^2^=0.48-0.63). Most of these models performed significantly better when considering genetic linkage between mutations (*r*^2^=0.53-0.67), pointing to epistasis being an important determinant in the evolution of SARS-CoV-2. Overall, our results highlight the predictive power of intrahost diversity data, and document the evolutionary forces shaping the fitness of mutations. Such insights offer potential to forecast the emergence of future variants and ultimately inform the design of vaccine targets.

## Introduction

Since its emergence in late 2019^1^, SARS-CoV-2, the agent of the COVID-19 pandemic, spread globally and caused over seven million deaths worldwide as of 21 July 2024 (https://ourworldindata.org/grapher/cumulative-covid-deaths-region). On 30^th^ January 2020, the World Health Organisation (WHO) declared the pandemic a Public Health Emergency of International Concern, which was then declared over on 5^th^ May 2023 following falling death rates, the deployment of effective vaccines, and increased immunity in the global population (https://www.who.int/emergencies/diseases/novel-coronavirus-2019/interactive-timeline).

Throughout the COVID-19 pandemic, SARS-CoV-2 has exhibited a repeated pattern of lineage replacement, where multiple waves of genetically distinct lineages, termed ‘variants’, emerged and replaced the previous dominant variant in circulation^2^. These variants, including Delta, Omicron, and BA.2.86 (an Omicron sub-lineage sometimes referred to as “Pirola”), carry constellations of co-occurring mutations, a phenomenon known as genetic linkage. Some of these mutations confer a fitness advantage to the virus such as increased transmissibility, replication or immune evasion^3^. Additionally, some linked mutations have been shown to act synergistically^4,5^, conferring greater fitness advantages than the sum of their effects in isolation. This process, termed synergistic epistasis, may explain some of the patterns of genetic linkage observed. The fitness differentials provided by these clusters of co-adapted mutations allow viruses carrying them to outcompete the preceding variant in circulation.

SARS-CoV-2 accumulates about 15 mutations per year with an overall evolutionary rate of ∼1×10^-3^ substitutions per site per year. The accrual of genetic diversity at the genome level is driven by multiple evolutionary forces at different levels. Within a single infection, genome replication errors and the action of host immune RNA editing may generate *de novo* mutations in the viral population that are subjected to within-host selective pressures. Together, these processes are estimated to result in the acquisition of 1.3×10^-6^ mutations per nucleotide per replication cycle^6^. The intrahost viral population expands and reaches its peak population size around 2-5 days after infection (coinciding with symptom onset)^7^. The viral population then undergoes a narrow transmission bottleneck where only a small fraction of the intrahost viral population is passed on to the next individual in the transmission chain, with estimates as low as one to eight virions per transmission event within households^8^. Selective forces may play an important role during transmission, favouring alleles that improve transmission and gradually purging those that do not. As a result of these processes, the genetic diversity observed within hosts is expected to be substantially higher than that observed at the interhost level. However, since SARS-CoV-2 typically causes acute infections that last about 10-15 days post symptom onset^7^, with samples often collected shortly after symptoms manifest, only a few intrahost mutations have been observed during this short window^8,9^.

The relative contributions of these evolutionary forces - acting both at the intrahost level and during transmission - on the mutations observed in SARS-CoV-2 lineages remain unclear. Given the short evolutionary timescale and the narrow population bottleneck during transmission, the effects of genetic drift may interfere with the action of natural selection. Neutral or weakly deleterious alleles may be transmitted before they are purged, allowing them to be observed at considerable frequencies amongst sampled genomes, whereas there may be a time lag before weakly advantageous mutations reach high frequencies. As such, we hypothesised that by analysing intrahost allele frequencies, we may be able to predict the relative success of mutations arising in the evolution of SARS-CoV-2. In addition, we hypothesised that the linkage patterns observed at the genomic level may be recapitulated by those found within intrahost viral populations. Addressing these questions may provide insights into the trajectory of mutations in SARS-CoV-2.

SARS-CoV-2 is arguably one of the best-studied viruses, owing to concerted, global and multidisciplinary research efforts over the COVID-19 pandemic. One important outcome is the remarkable amount of genomic data that has been generated. As of July 2024, nearly 15 million complete genomes and over seven million sequencing libraries of SARS-CoV-2 have been deposited on the data repositories, GISAID^10,11^ and NCBI Sequence Read Archive (SRA)^12^, respectively. These datasets and their accompanying metadata, including sample collection dates, sequence quality, lineage designations and viral mutations, provide an unprecedented ‘Big Data’ resource for testing this hypothesis and addressing other important questions relating to viral evolution.

In this study, we leverage large-scale SARS-CoV-2 genomic datasets to investigate the relationships between the genetic diversity observed within hosts and the relative fitness of SARS-CoV-2 mutations. We first provide a detailed overview of the landscape of mutations observed in SARS-CoV-2, exploring broad patterns in the physiochemical and phenotypic effects of all non-synonymous mutations observed over the COVID-19 pandemic. Using a curated dataset of 7862 quality-controlled sequencing libraries generated from SARS-CoV-2 infections, we then test the extent to which intrahost diversity, together with other genetic and phenotypic traits, can predict the relative success of SARS-CoV-2 mutations. These insights are valuable for understanding the evolutionary forces shaping SARS-CoV-2, and may offer scope to inform the selection of vaccine targets as well as responding to other current and future viral threats.

## Results

### The landscape of mutations in SARS-CoV-2

To explore the patterns of mutations that have arisen over the COVID-19 pandemic, we analysed the metadata provided by GISAID^10,11^ for the first submitted SARS-CoV-2 sequence up to the 8^th^ July 2024, comprising nearly 15 million complete SARS-CoV-2 genomes. Within this dataset, we identified a total of 101,484 single amino acid variants (SAVs). This represents 52% of the 194,660 SAVs that could theoretically be observed considering that SARS-CoV-2 coding regions encode 9733 amino acids, excluding start and stop codons. The number of SAVs observed over time increased rapidly in the initial stages of the pandemic though rapidly tapered off, with half of all theoretical SAVs already observed by December 2022 (**Fig. 1a**). Of all the SAVs observed in our dataset, 101,263 (99.8%) never exceeded a monthly global frequency of 10%, 100 (0.1%) reached frequencies between 10-90%, and 121 (0.1%) reached greater than 90% frequency (i.e., fixation). There were no obvious protein-specific patterns in the number of mutations observed for the vast majority of SAVs that failed to reached fixation (**Extended Data Fig. 1**). In contrast, SAVs that did were largely concentrated in the spike protein (**Extended Data Fig. 1**), consistent with its high antigenicity and crucial role in cellular entry.

**Figure 1.**
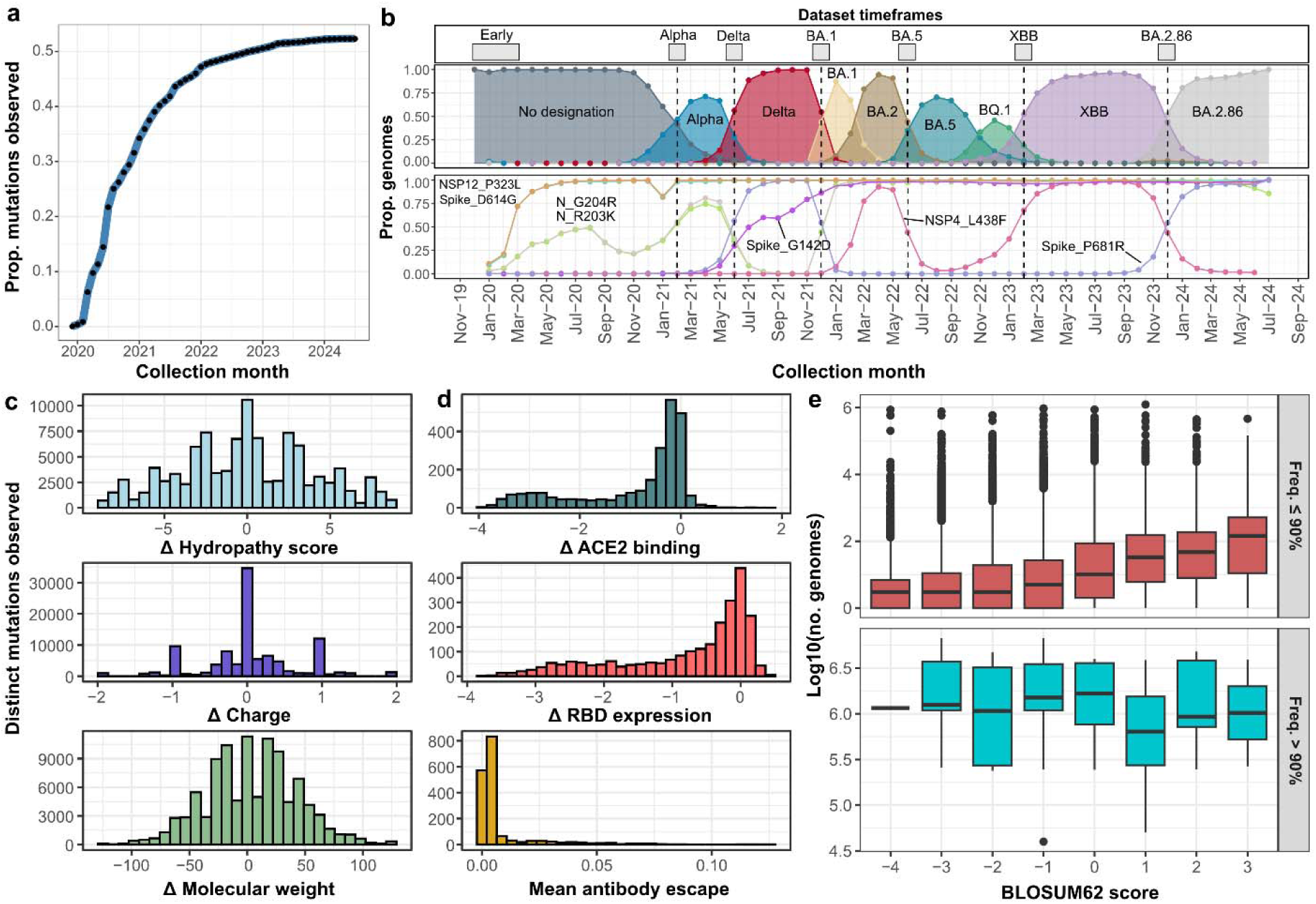
The phenotypic landscape of mutations in SARS-CoV-2. (a) Number of distinct single amino acid variants (SAVs) observed at various time points over the COVID-19 pandemic, normalised by the number of SAVs that could theoretically be observed (n=194,660). (b) Monthly frequencies of different lineages of interest (top) and a representative set of SAVs (bottom). Distributions of the (c) physiochemical effects and (d) phenotypic effects, as measured by deep mutational scanning (DMS) experiments^17,18^, of all SAVs observed in our dataset. (e) Boxplot of the log10-transformed genome count as a proxy for fitness stratified by BLOSUM62 score, for all SAVs that reached fixation (>90% monthly frequency) and those that did not.

The evolution of SARS-CoV-2 is largely characterised by the repeated emergence of divergent lineages that replace the preceding dominant lineage in circulation (**Fig. 1b**). Many mutations that reached fixation at some point over the course of the pandemic exhibited frequency patterns largely resembling those of these dominant lineages (**Fig. 1b**). However, there were some exceptions. For instance, Spike_D614G and NSP12_P323L rose in frequency during the initial stages of the pandemic, reaching fixation around mid-2020 and subsequently remained fixed in the SARS-CoV-2 population. Spike_G142D gradually rose in frequency during the Delta wave and remained fixed, while NSP4_L438F rose in frequency slightly after the initial Omicron lineages had replaced the preceding Delta variant and quickly dropped in frequency before reaching fixation again. This coincided with the emergence of the XBB/EG.5 Omicron sub-lineages (**Fig. 1b**).

We next explored whether the phenotypic impacts of SAVs observed were potential drivers of their relative fitness, which we estimate as the number of genomes submitted to GISAID carrying each SAV. This heuristic, which we henceforth refer to as ‘mutational fitness’ aims to approximate the ‘reproductive’ success of an individual mutation. It does not explicitly measure the contribution of a given mutation to the fitness of SARS-CoV-2, though the two are likely interlinked. Supporting analyses indicate that this metric is largely robust to sampling biases inherent to public sequence databases, and mostly recapitulates previously-derived fitness estimates^13^ (**Supplementary Note 1**).

We first calculated BLOSUM62 scores for all SAVs using the BLOSUM62 matrix, which evaluates amino acid changes based on their likelihood of occurring by chance, as derived from empirical protein alignments^14^. Under this scoring system, non-conservative mutations (*i.e.,* rare mutations) are assigned more negative scores^14,15^. While this scoring system is typically used for scoring amino acid sequence alignments, it is thought to capture the effects of differing selective forces on amino acid substitutions, where stronger purifying selection acts on less conservative mutations^15,16^. Since ‘conservativeness’ is a statistical abstraction of the phenotypic effects of mutation and the consequences of natural selection, we further considered other metrics that capture these effects directly. In particular, we computed the predicted effects of each SAV on the physiochemical properties of proteins (i.e., change in charge, molecular weight and hydropathy). We also investigated the effects of SAVs found within the receptor-binding domain (RBD) on protein expression, host-receptor binding and antibody escape by leveraging direct measurements from previously published Deep Mutational Scanning (DMS) experiments^4,17,18^ (see **Methods**).

Across SAVs, the score distributions for the three physiochemical properties (i.e., change in charge, molecular weight and hydropathy) appear symmetrically centred around zero (**Fig. 1c**), suggesting that most observed SAVs tend to preserve the physiochemical properties of the proteins they are found in. Separately, based on the DMS measurements, 83% and 70% of SAVs in the RBD were detrimental for host-receptor binding and protein expression, respectively, and most conferred minimal antibody escape (**Fig. 1d**). These results suggest that most SAVs observed are neutral or deleterious for these phenotypes.

Notably, we found a significant positive correlation between fitness and the BLOSUM62 score of all SAVs (Spearman’s ρ=0.33, *p*<0.0001; **Fig. 1e**), consistent with the idea that non-conservative mutations are generally deleterious^19–21^. This relationship, however, does not hold for the small subset of SAVs that have reached fixation over the course of the pandemic (ρ=-0.068, *p*=0.462; **Fig. 1e**), which are likely to confer the highest fitness gains. These findings suggest that while non-conservative mutations are generally deleterious, they are also more likely to confer the highest fitness gains to the virus. This is further corroborated by the relationships observed between the physiochemical effects of SAVs and fitness. Indeed, the absolute changes in physiochemical scores were significantly and negatively correlated with fitness (ρ=-0.040,-0.13,-0.23, respectively; all *p*<0.0001). Meanwhile, SAVs that reached fixation were associated with significantly higher absolute changes in charge compared to non-fixed SAVs (two-sided Mann-Whitney U test, p=0.0076). These results highlight that while more significant biochemical alterations are often deleterious, they can occasionally lead to the highest fitness gains.

Finally, SAVs that are associated with increased host-receptor binding and RBD expression had significantly higher fitness values (both ρ=0.26, p<0.0001), which is expected given the key role of the RBD in mediating viral entry into host cells. Meanwhile, antibody escape was weakly and negatively correlated with fitness (ρ=-0.084, p=0.0005). While SAVs that improve antibody escape would likely be beneficial for the virus, and therefore be inherently fitter, this negative correlation may point to a fitness trade-off with other phenotypes. Indeed, antibody escape was significantly negatively correlated with both binding (ρ=-0.36, p<0.0001) and expression (ρ=-0.11, p<0.0001). This indicates that the fitness advantage associated with increased antibody escape may be counteracted by decreased binding and/or expression, thus resulting in a lower overall fitness, a phenomenon noted previously^22^. However, there are some exceptions to this trade-off such as the VoC-associated RBD SAVs, Spike_L452R and Spike_E484K, whose effects on binding and antibody escape are both larger than the 90th percentile of associated DMS measurements.

### Intrahost dynamics predicts the future fitness of mutations

We next investigated whether the intrahost diversity of SARS-CoV-2 infections could be used to forecast the relative fitness of SAVs at the population level. To do this, we curated and downloaded seven sequencing datasets (total n=7862) comprising sequencing libraries of SARS-CoV-2 infections, which were sampled at various timepoints across the pandemic. These datasets correspond to seven time periods of sampling, henceforth timeframes, termed ‘Early’, ‘Alpha’, ‘Delta’, ‘BA.1, ‘BA.5’, ‘XBB’ and ‘BA.2.86’ (**Fig. 1b**). The Early timeframe corresponds to the first four months of the pandemic, while the others span the transition points between the major variant waves in our dataset. We estimated the intrahost frequencies of SAVs in these libraries using stringent filtering criteria (see **Methods**)^8^. In contrast to previous intrahost diversity studies that focused only on subconsensus SAVs (<50% intrahost frequency)^8,23–25^, we additionally considered the consensus SAVs (≥50% intrahost frequency) in our samples, as both SAV subsets reflect intrahost selective pressures.

To ensure reliable estimation of intrahost diversity, we applied quality control filters to remove libraries potentially associated with serial passaging experiments, laboratory contamination, an excess of sequencing artefacts, or erroneous collection dates (see **Methods** and **Supplementary Note 2**). Additionally, previous studies have noted that cross-contamination of samples may result in the detection of erroneous intrahost SAVs^8,23^. While we could not completely rule out the possibility of cross-contamination, our analyses indicate that cross-contamination is not a major confounder in our study (**Supplementary Note 3**). Our final quality-controlled dataset comprised 5870 sequencing libraries, representing seven snapshots of the lineage composition at various points during the COVID-19 pandemic (**Extended Data Fig. 2**).

Within each of the seven datasets, we identified between 2927-11,992 intrahost SAVs across the sequencing libraries, with an increase in the median number of SAVs detected per library by the chronological order of the datasets (Early=24; BA.2.86=98). Within each timeframe, 9-110 (0.10-3.8%) of these intrahost SAVs were already at >10% monthly frequency at the interhost level. Excluding these SAVs, we found that 5-75 (0.18-1.1%) SAVs reached at least a monthly frequency of >10% after the timeframe of each dataset. Further, these intrahost SAVs only reached >10% population frequency at a median of 3-22 months after the timeframe of each dataset (**Fig. 2a**). In other words, many of these relatively fitter SAVs could already be observed in the intrahost diversity of infections that were sampled much earlier in the pandemic. Analysis of an independent intrahost dataset^23^ (henceforth ‘Tonkin-Hill dataset’) – specifically generated for studying the intrahost diversity of SARS-CoV-2 infections – found that many intrahost SAVs reached >10% frequency at a median of 19 months after the timeframe of the dataset (**Supplementary Note 3**), further supporting our observations.

**Figure 2.**
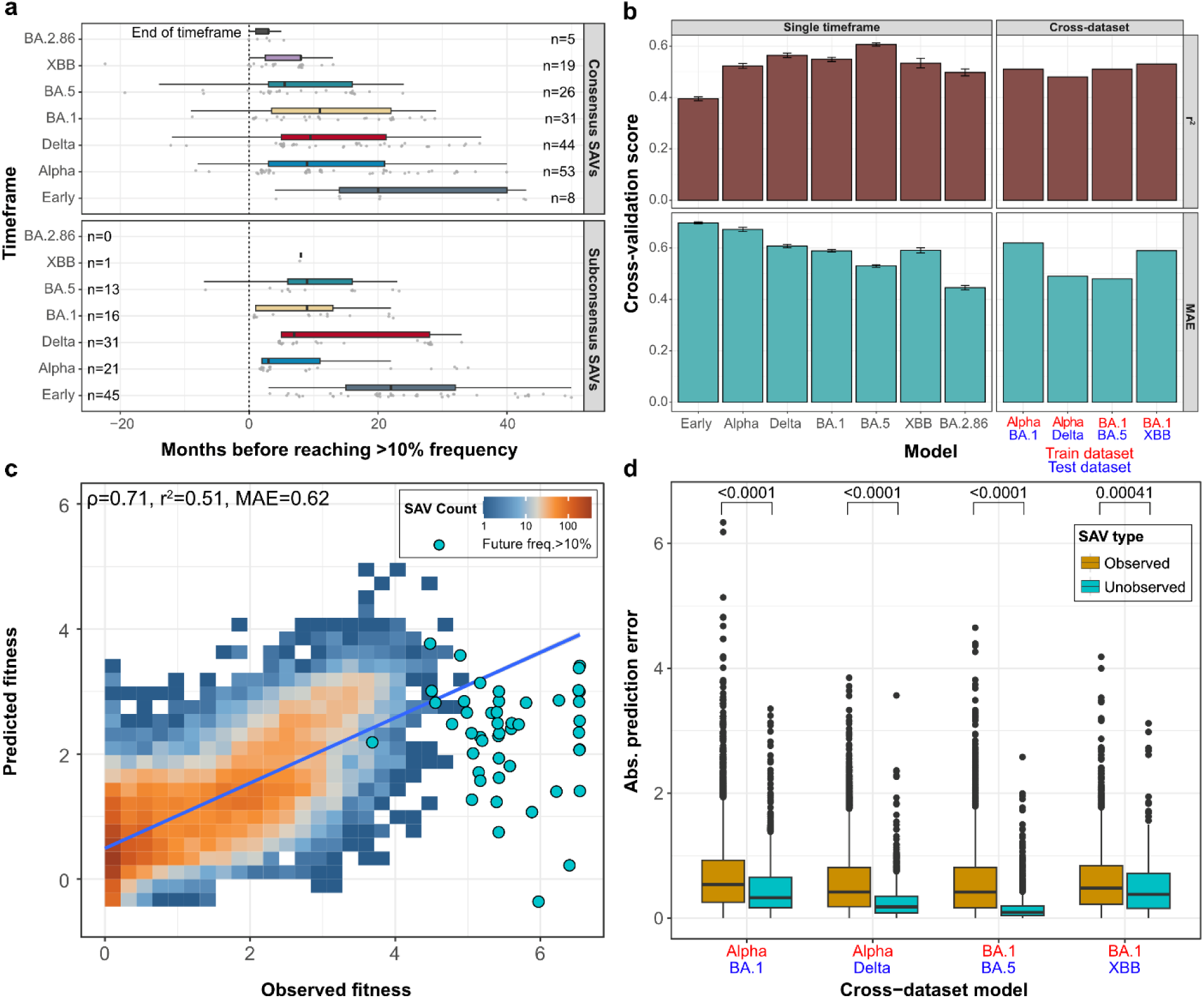
Intrahost diversity predicts the success of mutations. (a) Time difference between the end of each timeframe and the first month where an intrahost SAV reaches >10% at the population frequency. Data is stratified by subconsensus and consensus SAVs (intrahost frequency ≥50% and <50%, respectively). The number of intrahost SAVs found in each dataset is annotated. (b) Prediction performance of our single timeframe and cross-dataset models. The former set of models were evaluated using a nested cross-validation procedure and the latter by training models on intrahost data from one timeframe and testing on data from another. Error bars for single timeframe models represent the standard error of scores across all cross-validation splits. (c) Representative density plot showing the high correlation between the predicted future fitness and the actual observed fitness of SAVs in our cross-dataset models. Data shown is from a model trained on the Alpha timeframe and evaluated on the BA.1 timeframe. SAVs that reached a high future frequency (i.e., >10%) are shown as discrete points. The blue line indicates a linear regression smooth of the predicted and observed SAV fitness values. (d) Absolute prediction errors of SAVs that were not observed during or before the end of the test timeframe, and those that were, for our cross-dataset models. Differences in distributions were tested using two-sided Mann-Whitney U tests and the corresponding p-values are annotated. Boxplot elements are defined as follows: centre line, median; box limits, upper and lower quartiles; whiskers, 1.5x interquartile range. In (b) and (d), r^2^ refers to the coefficient of determination score for regression models.

Next, we trained supervised XGBoost^26^ regression models to forecast the fitness of each intrahost SAV after the timeframe of each dataset (henceforth ‘future fitness’), which we estimated as the number of submitted consensus genomes a SAV was observed in. These models leveraged 13 predictors of future fitness, capturing the intrahost dynamics, and physiochemical and phenotypic effects of each SAV. These were the number of libraries each SAV was detected in, median and maximum intrahost frequency across all samples, BLOSUM62 score, the estimated raw and absolute change in charge, hydropathy and molecular weight, and for RBD mutations, the change in binding and expression, and antibody escape fraction, as directly measured via DMS^4,17,18^. Most predictors were significantly correlated with the future fitness of SAVs (Spearman’s correlation test, adjusted p<0.05) and the direction of the correlations were largely consistent across datasets (**Extended Data Fig. 3**). Maximum and median intrahost frequency, BLOSUM62 score, hydrophobicity and molecular weight were positively correlated with future fitness, whereas absolute changes to charge, molecular weight and hydropathy were negatively correlated with future fitness (**Extended Data Fig. 3**). For each timeframe, we optimised a fitness model and assessed its performance using a nested cross-validation procedure (see **Methods**), where different subsets of the intrahost data are withheld to test whether the model can make robust predictions on unseen data. Across all timeframes, models were able to predict the future fitness of intrahost SAVs, yielding mean *r*^2^ values between 48-63% (**Fig. 2b**). Additionally, the mean absolute error of the predictions by each model was 0.45-0.68 (**Fig. 2b**), indicating that, on average, the models were able to predict the future fitness of intrahost SAVs to within the same order of magnitude. These findings indicate that our models could reliably forecast the success of intrahost SAVs using a combination of intrahost, physiochemical and phenotypic predictors.

To further test the generalisability of our models, we trained four models on data from specific timeframes, and assessed their performance on those from more recent timeframes (henceforth ‘cross-dataset models’). As an example, we trained a model on the Alpha dataset, comprising libraries collected in February 2021, to predict the future fitness of intrahost SAVs up to before the first Omicron wave (i.e., BA.1; November 2021). We then tested the trained model on the BA.1 dataset (libraries collected in December 2021), to predict the fitness of intrahost SAVs from BA.1 to the end of our dataset (July 2024) (**Fig. 2c**). Overall, all four cross-dataset models were able to reliably predict mutational fitness on an independent dataset (Spearman’s ρ=0.66-0.72, r^2^=0.52-0.60; **Fig. 2b**), indicating that our models are able to generalise across intrahost data from different timeframes. Additionally, for these models, the prediction errors for SAVs in different protein regions were comparable (**Extended Data Fig. 4**), suggesting the models are able to generalise across different proteins despite their varied functions and antigenicity. Furthermore, two other cross-dataset models that were trained and evaluated on data from different geographical regions showed comparable results (ρ=0.70 and 0.68, respectively; both r^2^=0.50). This further highlights the robustness of our modelling framework to sampling biases (**Supplementary Note 1**).

We then tested the predictive power of intrahost data, specifically, whether our models can reliably predict the future fitness of intrahost SAVs that have not been identified at the population level before or during the timeframe of each dataset. To do this, we performed post-hoc analyses on the cross-dataset models, comparing the absolute prediction errors of previously ‘unobserved’ SAVs to those already observed by the end of the test timeframe. For all models, the prediction errors for the ‘unobserved’ mutations were significantly lower than for the ‘observed’ mutations (Mann-Whitney U test, all p<0.0001; **Fig. 2d**). These results indicate that our models can in fact predict the future success of unobserved mutations, highlighting the predictive power of viral intrahost diversity.

Notably, while some of the intrahost SAVs that reach a future frequency >10% are predicted by our cross-dataset models to have the highest future fitness, many of the high frequency SAVs had markedly lower predicted fitness values than observed (**Fig. 2c**). Indeed, the studentised residuals for high frequency SAVs (>10% future fitness) – in a linear model of predicted versus observed fitness – were considerably more negative than those for low frequency SAVs (**Extended Data Fig. 5**). This prediction error may be a result of other evolutionary processes not captured by the model including epistatic interactions between high frequency mutations.

### The biological correlates of fitness

To understand how these complex models inform their predictions, we further used the model interpretation framework, SHAP^27^, to visualise the relationships between the predictors and mutation fitness. The core of this framework is the SHAP value, which here represents, for each SAV, the estimated contribution of a single predictor to the predicted fitness value for that SAV. Across the datasets, the median absolute SHAP value for maximum intrahost frequency was the highest, indicating that this predictor was the most important contributor to the model predictions (**Fig. 3a**). This further suggests that success at the intrahost level is a strong correlate of the relative fitness of SAVs at the population level. However, models leveraging only intrahost predictors did not perform as well as our full models comprising all 13 predictors (**Extended Data Fig. 6**), suggesting that the physiochemical and phenotypic predictors are also key contributors to predicting the future fitness of intrahost SAVs.

**Figure 3.**
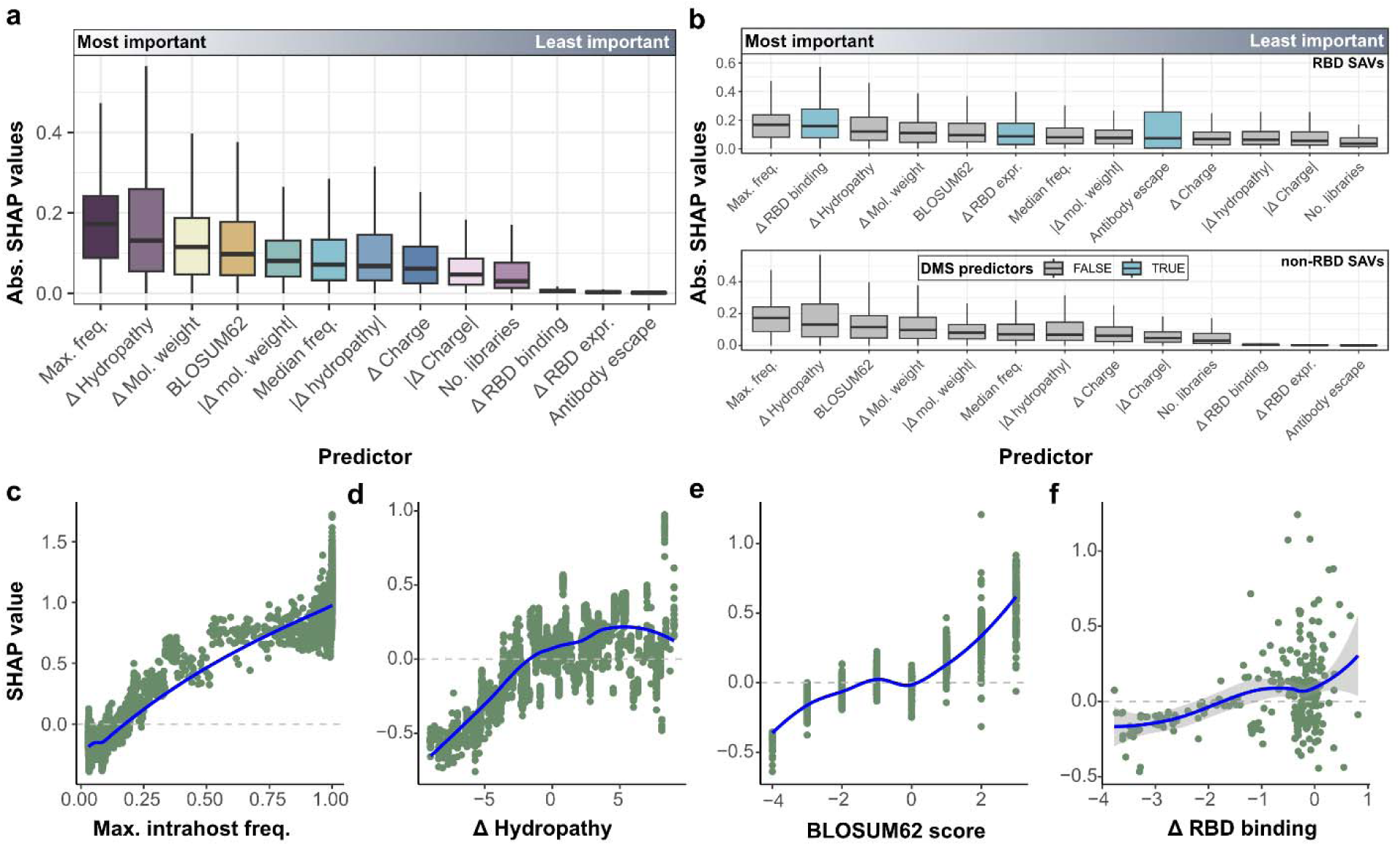
Models are able to capture biologically meaningful correlates of fitness. (a) Relative importance of predictors for all fitness models trained, as assessed using SHAP values. Higher absolute SHAP values indicate higher contributions to the model predictions. Boxplots were ranked by the median absolute SHAP value of each predictor. (b) Boxplots showing the relative importance of predictors for all fitness models, as in (a), but stratified by RBD and non-RBD SAVs. For (a) and (b), predictors are ranked from most important (highest median absolute SHAP value) to least important. Scatter plots showing the relationships between future mutational fitness predicted by our XGBoost regression models and the (c) maximum intrahost frequency, (d) change in hydropathy, (e) BLOSUM62 score, and (f) change in RBD binding for each SAV. The data shown in (c)-(f) were generated using the Delta dataset, but are representative of the relationships observed in the other datasets considered. Blue lines represent LOESS (Locally Estimated Scatterplot Smoothing) regression smooths for each scatter plot.

Mutation-by-mutation DMS measurements of the phenotypic impact on RBD binding, expression, and mean antibody escape were the least important contributors overall (**Fig. 3a**). However, we suggest that this is likely due to the fact that these phenotypic predictors are only available for RBD SAVs and hence are not informative for predicting the fitness of non-RBD SAVs. Stratifying by mutation type, we found that all three phenotypic predictors are much more important for predicting the future fitness of RBD SAVs than for non-RBD SAVs (**Fig. 3b**). This suggests that our models are also able to capture the functional and antigenic importance of the RBD.

We additionally explored the relationships between each of the predictors and the future fitness of SAVs by evaluating their corresponding SHAP values (**Fig. 3c-f**). Maximum intrahost frequency had a monotonic positive relationship with predicted future fitness (**Fig. 3c**), indicating that the selective forces governing the intrahost frequencies of SAVs explain, in part, mutational success at the global level. Separately, there was an apparent parabolic relationship between fitness and change in hydropathy, where SAVs involving relatively larger increases or decreases to hydrophobicity are generally associated with lower mutational fitness (**Fig. 3d**). Similarly, BLOSUM62 score was positively correlated with predicted fitness (**Fig. 3e**). These findings recapitulate the previously observed pattern (**Fig. 1c-e**) that more marked physiochemical alterations are typically associated with lower fitness. However, we note that some SAVs associated with larger physiochemical changes had the most positive SHAP values (**Extended Data Fig. 7**). This suggests that whilst large physiochemical changes are generally deleterious, they can also lead to positive fitness gains. A separate analysis of SAVs exclusively in the GISAID dataset indicated that high-fitness SAVs estimated to cause large physiochemical changes are enriched in the spike protein. Such mutations may potentially be linked to the SARS-CoV-2 spike protein adapting to onward transmission in humans (**Supplementary Note 4**). Finally, RBD binding (**Fig. 3f**) and expression (**Extended Data Fig. 7**) were both positively correlated with higher predicted fitness values. These findings indicate that, as expected, SAVs associated with increases in binding and expression are fitter. All patterns were largely concordant across the different timeframes considered (**Extended Data Fig. 7**), highlighting that our models learned biologically relevant patterns to predict the future fitness of mutations.

### Intrahost diversity recapitulates interhost linkage patterns

We next explored whether genetic linkage at either the intrahost or interhost level could explain why some of these SAVs are much fitter than predicted by our initial models (**Fig. 2c**; **Extended Data Fig. 5**). To estimate intrahost linkage, we calculated Pearson’s *r* using the intrahost SAV frequencies within each dataset. At the interhost level, we calculated the *D’* statistic^28^ using the SAV frequencies across the entirety of GISAID submissions. To reduce noise arising from low frequency SAVs, we considered only SAVs that were observed in more than 1000 genomes. For biallelic loci (i.e., only two alleles or amino acid variants are observed at each locus), linkage is often reported as *|D’|* or *r*^2^ since negative values indicate linkage between the alternate allele at one locus and the wild type at the other. In our context, there are 20 possible SAVs at each locus so negative *D’* or *r* values are not as meaningful and can only be interpreted as ‘not positively linked’.

In total, we calculated *D’* for 9,100,120 SAV pairs across the entire GISAID dataset, and *r* for 96,945-1,972,800 SAV pairs for each intrahost dataset, of which 0.19% and 0.01-1.7% were strongly linked (*D’* or *r*>0.9), respectively. At the interhost level, within-protein linkage associations were significantly more common in strongly and positively linked (*D’*>0.9) SAVs than those with weaker linkage associations (*D’*≤0.9) (**Fig. 4a**; odds ratio=3.79, Fisher’s exact test, p<0.0001). This is consistent with the expectation that linkage decays with physical distance, since recombination is more likely to break up associations between more distant loci and that short-range epistatic interactions are expected to be more common. We found a similar enrichment of strong within-protein linkage patterns across all intrahost datasets (**Fig. 4a**; odds ratio=1.7-44, all p<0.0001), indicating that intrahost linkage patterns also capture the effects of these evolutionary mechanisms.

**Figure 4.**
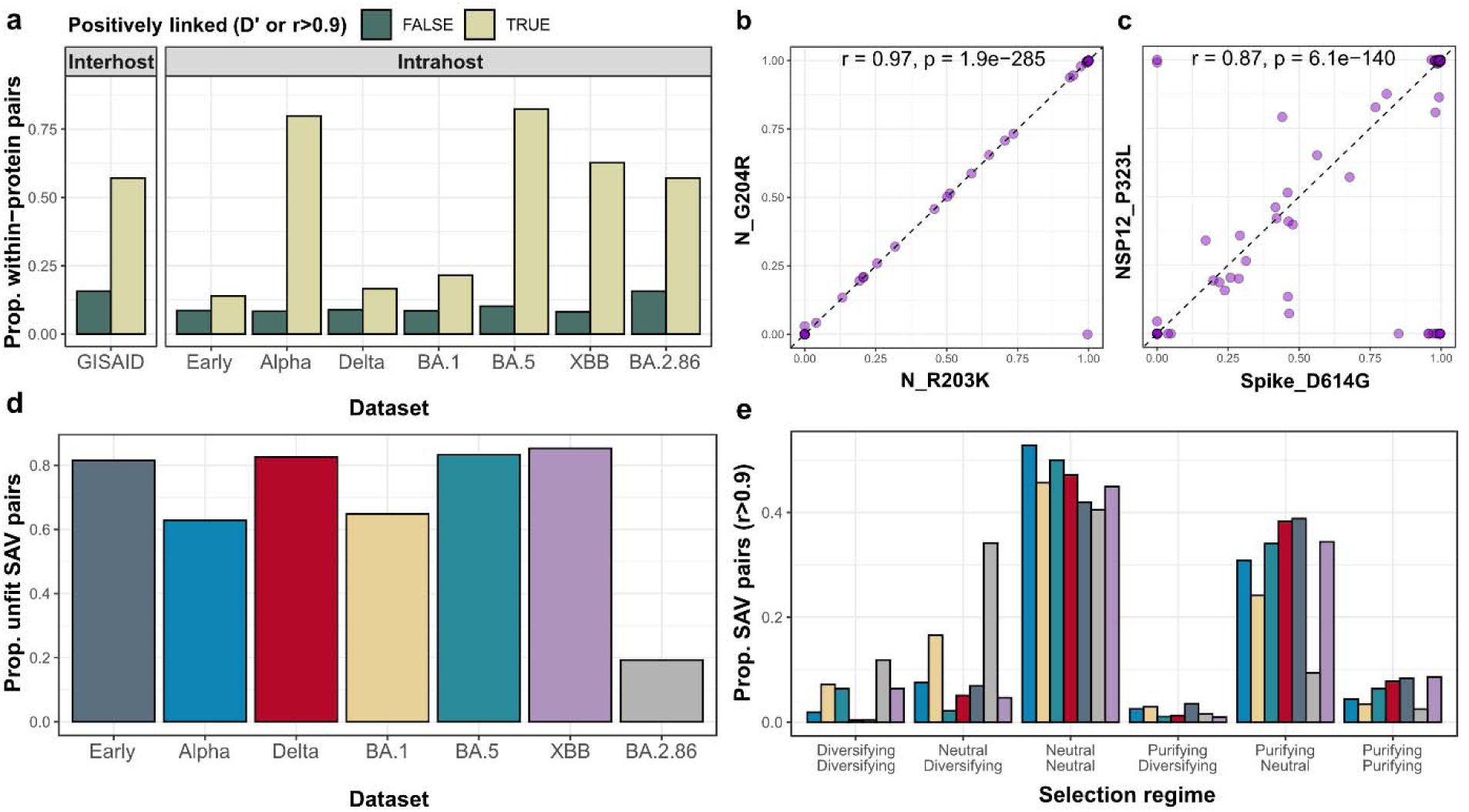
Intrahost evolution shapes global linkage patterns. (a) Proportion of SAV pairs involving SAVs that are found within the same protein at the (top) interhost or (bottom) intrahost level. Scatter plots showing the correlations between the intrahost frequencies of (b) N_R203K and N_G204R and (c) Spike_D614G and NSP12_P323L. (d) Proportion of strongly-linked SAV pairs (r>0.9) that only involve low fitness SAVs, defined as those that were observed in at most 1000 consensus genomes across the 56 months. (e) Proportion of strongly-linked SAV pairs (r>0.9) associated with different combinations of selection regimes. Each codon associated with an SAV were classified as under positive, negative or neutral selection based on selection analysis data produced by Sergei Pond (https://observablehq.com/@spond/sars-cov-2-global-genomic-selection-2019-aug-2023). This data was generated using the Fixed Effects Likelihood (FEL) method^32^ in HyPhy^31^.

The SAV pairs that appear strongly linked at both the intrahost and interhost level included SAVs that were highly successful, including N_R203K - N_G204R (*r*=0.98 and D’=1.00 for the Alpha timeframe; **Fig. 4b**). The close proximity of these two SAVs and the near-perfect linkage patterns indicate likely synergistic interactions. Separately, Spike_D614G - NSP12_P323L, one of the first pairs of SAVs whose monthly frequencies co-rose rapidly in the early stages of the pandemic (**Fig. 1b**), had a slightly weaker intrahost correlation (‘Early’ timeframe, intrahost *r*=0.84; **Fig. 4b**). The occurrence of these two SAVs on different proteins and the weaker linkage pattern may point to co-selection of two highly beneficial mutations^29,30^ in a largely wild-type background, rather than epistasis.

Notably, across datasets, a large proportion (19-85%) of strongly linked pairs (r>0.9) involved only SAVs that were observed in at most 1000 consensus genomes across the 56 months (**Fig. 4d**). The low consensus prevalence of these SAVs indicate that the majority of linked SAVs do not confer a significant fitness advantage and are likely a product of genetic drift. Concordantly, a formal analysis of the differential selective regimes that SAVs are evolving under^31,32^ (**Supplementary Note 5**) indicates that the majority of strongly linked SAV pairs (r>0.9) comprised SAVs that were inferred to be either under neutral and neutral, or purifying and neutral selection, respectively (**Fig. 4e**). The former and latter categories potentially represent linkage patterns that may derive from genetic drift, or compensatory epistasis, respectively. However, disentangling the exact evolutionary mechanisms underlying these patterns require further experimental work, as exemplified by previous studies^4,5,33^.

Overall, our findings indicate that the linkage patterns observed at the interhost level can also be seen at the intrahost level. However, our intrahost linkage measures are more noisy in that they capture linkage patterns that may not be associated with mutational fitness.

### Genetic linkage improves fitness predictions

To incorporate genetic linkage at both the intrahost and interhost levels into our models, we generated six further predictors, in addition to the thirteen used previously, namely, whether the SAV is strongly linked to at least one other SAV (*D’* or *r*>0.9), the number of strongly linked SAVs (*D’* or *r*>0.9), and the maximum *D’* or *r* across all pairs involving the SAV. Here, *D’* statistics were only calculated for SAVs observed in >1000 consensus genomes *prior* to the end of each timeframe. However, given that prior fitness (i.e., presence in >1000 consensus genomes) is highly autocorrelated with future fitness (Spearman’s ρ=0.88-0.92), models may ‘cheat’ by leveraging this autocorrelation to improve performance without learning any additional biological patterns. As such, we included an additional binary predictor, indicating whether a SAV was observed in >1000 genomes (henceforth ‘prior fitness’), to partition this effect in our models.

Inclusion of linkage predictors significantly improved the predictive performance of all models (*r*^2^=0.59-0.67; Mann-Whitney U test, maximum *p*=0.029), except for that trained on the Alpha timeframe (*r*^2^=0.53; *p*=0.436) (**Fig. 5a**). The most prominent improvement was seen for the model trained on the BA.2.86 timeframe (**Fig. 5a**). All cross-dataset models trained on one timeframe and tested on another (as for **Fig. 2b-d**) also performed better with the inclusion of linkage predictors (**Extended Data Fig. 8**). This indicates that the relationships between linkage and future fitness learned by the regression models were generalisable across datasets. Since SHAP values are additive^27^, we compared the sum of the absolute SHAP values for the different types of predictors (i.e., physiochemical, phenotypic, intrahost, intrahost linkage, interhost linkage and prior fitness) to assess their relative importance. Interhost linkage predictors were significantly more important than intrahost linkage predictors (paired *t*-test, *t*=22.6-107, d.f.=2816-11935, all p<0.0001) and the prior fitness predictor for predicting future fitness, across all timeframes (*t*=30.9-114, d.f.=2761-9560, all p<0.0001) (**Fig. 5b**). These findings indicate that linkage patterns at the interhost level are largely responsible for the performance boost. This is likely because our interhost predictors, which rely on linkage patterns in consensus genomes, are better at capturing allele associations driven by epistatic selection, whereas intrahost linkage associations tend to include a larger fraction of randomly co-occurring SAVs (**Fig. 4d** and **e**; **Supplementary Note 5**).

**Figure 5.**
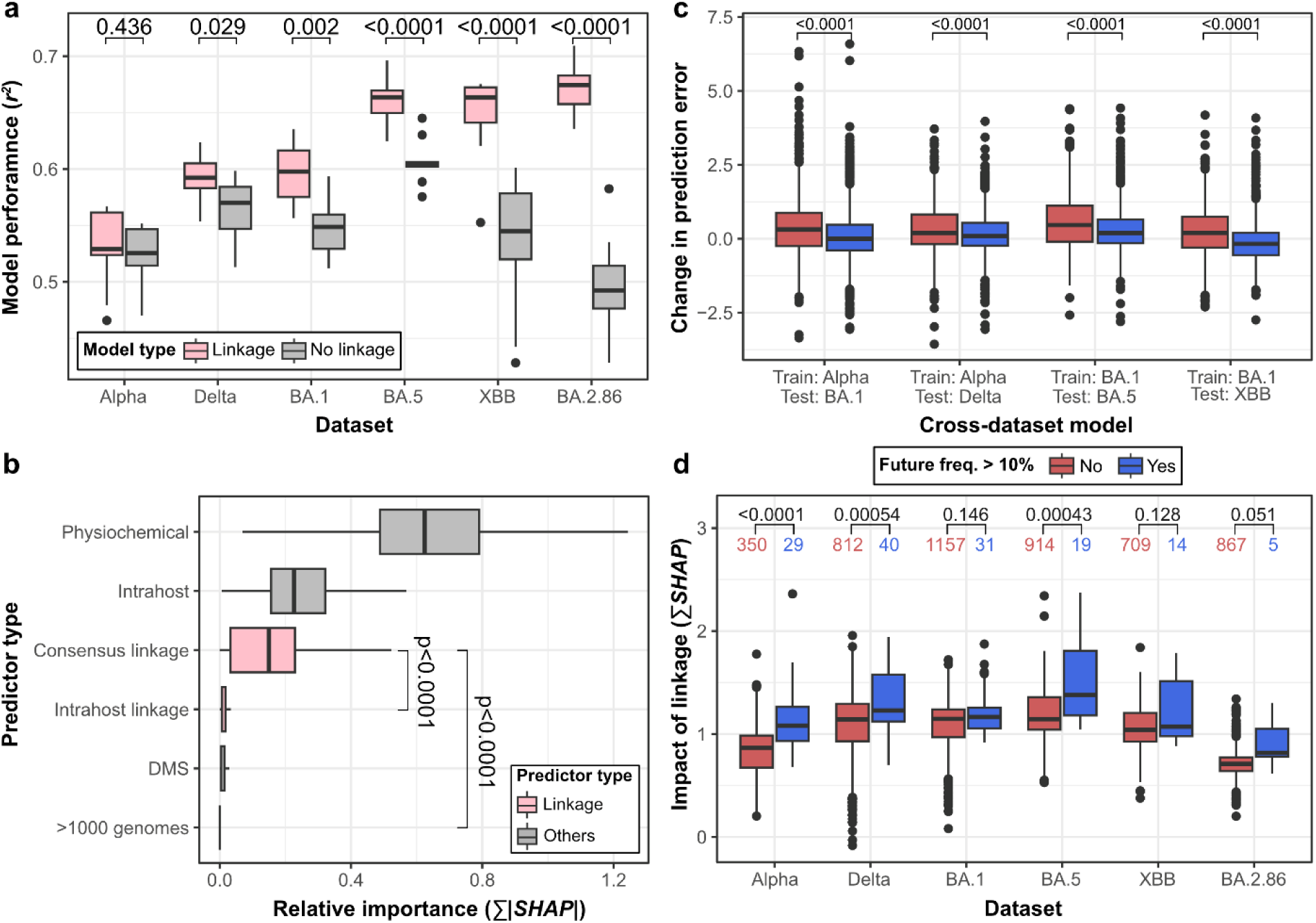
Consideration of genetic linkage improves fitness predictions. (a) Nested cross-validation performance (assessed via the coefficient of determination score, *r^2^*) of single timeframe models with and without the inclusion of linkage predictors. (b) Distributions of the summed absolute SHAP values for each predictor category reflecting the aggregated impact of each predictor type. Predictor types are ranked from top to bottom by decreasing importance, as determined via their median absolute SHAP values. Differences in the relative importance of each predictor category were tested using two-sided paired *t*-tests for each dataset. The tests for each comparison and for each dataset were highly significant (all p<0.0001). (c) The changes in prediction error (i.e., |*ε_linkage_*| - |*ε_no linkage_*|) for the Alpha, Delta, BA.1, BA.5, XBB and BA.2.86 datasets after including linkage predictors, stratified by whether a SAV was observed at >10% monthly frequency after the timeframe of the dataset. (d) Impact of linkage predictors (i.e., ∑(***SHAP***)) on the predicted fitness of SAVs, stratified in the same way. Boxplot elements are defined as follows: centre line, median; box limits, upper and lower quartiles; whiskers, 1.5x interquartile range. Differences in distributions were tested using two-sided Mann-Whitney U tests and the corresponding p-values are annotated.

Since the fittest SAVs were associated with significantly higher prediction errors than the less successful SAVs in our initial models (**Fig. 2c**; **Extended Data Fig. 5**), we hypothesised that genetic linkage may explain, at least in part, this pattern. To test this, we compared the changes in prediction error for our models with or without linkage predictors for SAVs where *D’* could be calculated. Following the inclusion of linkage predictors, the changes in prediction errors were significantly more negative for fitter SAVs (future frequency>10%) compared to less successful SAVs (Mann-Whitney U test, all p<0.001, **Fig. 5c**). Additionally, we aggregated the contributions of all linkage predictors by summing their SHAP values and found more positive SHAP values for the fitter SAVs (future frequency>10%) (**Fig. 5d**). This suggests that the predicted fitness scores of the fitter SAVs were increased to a larger extent than for less fit mutations (future frequency≤10%) due to the addition of linkage predictors. These results jointly indicate that by using information that captures genetic linkage, these models were able to predict the higher future fitness values for the fitter SAVs, thus resulting in lower overall prediction errors. Together, this suggests that the success of high frequency SAVs may, in part, be governed by epistatic interactions.

## Discussion

Predicting which mutations will emerge and reach high frequency in the future is a long-standing and important, yet largely unsolved problem in the study of pathogen dynamics. In this study, we investigated whether the intrahost diversity of SARS-CoV-2 infections could provide clues on the relative fitness of SAVs by leveraging the unparalleled amount of sequencing data that has been generated and shared. Our key assumption is that the evolutionary forces shaping the interhost frequencies of mutations in the viral population are also acting within hosts. Using various machine learning techniques, we demonstrated that this assumption holds, that is, that the intrahost dynamics of SARS-CoV-2 infections can well predict the future fitness of mutations. In addition, our model interpretation analyses revealed highly generalisable patterns of intrahost, physiochemical and phenotypic traits that correlate with the success of mutations. Our findings lay the groundwork for developing better predictive tools that may be invaluable for pre-empting the emergence of novel lineages and supporting the design of vaccine targets.

By analysing the distribution of SAVs across all complete genomes on GISAID (as of July 2024), we found that more than half of all possible SAVs have already emerged in the evolution of SARS-CoV-2, and that the rate of emergence of previously unseen mutations has slowed (Fig. 1a). Most SAVs observed thus far tend to preserve the physiochemical properties of the proteins they are found in, suggesting that major phenotypic alterations to proteins are generally deleterious (Fig. 1c). This is corroborated by the observation that most SAVs in the RBD are estimated to be phenotypically neutral or deleterious (Fig. 1d). Further, only an extremely small subset of SAVs have reached a high frequency at the interhost level thus far. Collectively, these findings point to strong functional and evolutionary constraints on the mutational space that SARS-CoV-2 can explore. Since the rate of emergence of mutations has slowed considerably, one can speculate that future lineages may be more likely to harbour novel combinations of moderately or highly fit mutations that have already been observed so far rather than a set of previously unseen mutations. This, however, does not preclude the possibility of unseen mutations emerging in the future.

In this study, we used the number of GISAID sequences carrying each SAV across various time windows as a proxy for estimating the ‘reproductive’ success of SAVs, which we refer to as ‘mutational fitness’. Our models are therefore trained to predict how successful a mutation is rather than the contribution of a mutation to the fitness of individual viral lineages. In comparison, previous fitness metrics, such as those developed by Bloom and Neher^13^, and by Donker and colleagues^34^ estimate the relative fitness contribution of mutations to SARS-CoV-2 lineages using GISAID data. The two approaches are tightly interlinked, and indeed our fitness scores are highly correlated (Spearman’s ρ=0.73) with the Bloom and Neher’s^13^ fitness effects scores. One important distinction is that these more derived metrics^13,34^ impose priors on the underlying model of fitness effects. Bloom and Neher’s^13^ measurement of fitness effects is based on the number of independent emergences of mutations along a tree, so it is heavily reliant on robust phylogenetic reconstruction and minimal recombination. Donker et al.^34^ estimate fitness effects based on the speed at which a novel mutation replaces others across two timepoints, which may be more sensitive to sampling biases. Meanwhile, our fitness metric does not impose any model priors on the data and may therefore be more generalisable and scalable.

That being said, all such fitness estimates (including ours) are likely subject to spatiotemporal biases in genomic surveillance efforts^35^. For example, the estimated fitness of a moderately successful SAV that circulated only in countries with lower surveillance would be lower than a less successful SAV that spread in countries with more extensive surveillance. While we do not explicitly correct for these biases, we found through a cross-regional analysis that they do not significantly impact our fitness estimates (**Supplementary Note 1**). This is consistent with Bloom and Neher’s observation that their fitness effect scores are largely correlated across different geographical subsets of GISAID data^13^. Further, we could reproduce our findings using an independent dataset designed solely for the study of intrahost diversity^23^ (**Supplementary Note 1**), further highlighting the robustness of our results to sampling biases.

Our modelling approach differs from previous efforts to understand viral fitness, which have largely focused on the relative fitness of lineages^36–41^, whereas we consider individual mutations as targets of selection, each associated with its own fitness values. The theoretical number of combinations of SAVs that SARS-CoV-2 can have is practically infinite (i.e., 20^9733^), which entails that the fitness of previously unseen lineages is difficult to predict, especially for models based on epidemiological data^36,38^. Focusing on the fitness of mutations is lineage agnostic (i.e., not informed by epidemiological information about lineage circulation or frequency) and enables us to derive the inherent traits that allow SAVs to rise to high frequencies. For example, our models recapitulate previous findings^19–21^ that the SAVs associated with more marked physiochemical changes are generally under stronger purifying selection, and expectedly, that the SAVs that negatively impact RBD binding and expression are less fit. Further, our mutation-specific modelling framework can also be extended to infer the effects of contextual factors that are known to govern mutational success, including the current immune landscape (e.g. whether a mutation falls within T-cell epitopes, and the frequency of HLA types in the population that recognise these epitopes), infection histories (i.e., whether and when an SAV was previously circulating in the population), and the diversity of extant competing lineages in circulation (e.g. prior and current frequencies of competing SAVs at each locus).

Importantly, our modelling approach leverages information about the intrahost dynamics of mutations, viewing each infection as an independent playing field where mutations compete for dominance, with fitter mutations being observed more frequently than less fit ones. Our observation that maximum intrahost frequency is consistently the most important predictor of future fitness supports this, and suggests that within-host evolutionary processes are important components that shape the fitness landscape of mutations at the interhost level. Such processes may include positive selection for mutations that improve viral infectivity and/or replication rate, and purifying selection acting on highly deleterious mutations that drastically alter protein function. Separately, our results indicate that highly successful SAVs could be detected in the intrahost diversity of infections sampled long before their rise to success (Fig. 2a). This considerable ‘time lag’ may ultimately facilitate pre-emptive mitigation strategies including vaccine design.

Notably, our models performed relatively poorly on making predictions for the fittest mutations (Fig. 2c; **Extended Data** Fig. 5). Incorporating genetic linkage into the models partially reduces this effect, indicating that the co-occurrence of SAVs, which may or may not involve epistasis, plays a role in modulating the fitness of mutations in the evolution of SARS-CoV-2. However, there remains considerable variation in the fitness of SAVs that is not captured by our models, suggesting that some components of mutational fitness are still unaccounted for. Incorporation of further functional, structural or immunological predictors may reduce this gap further, but this may be challenging since existing experimental data for SARS-CoV-2 remain biased towards the spike protein. Additionally, selective forces may act at the point of transmission, with mutations improving transmissibility being more successful, but this is not explicitly captured by our models. Alternatively, these discrepancies may indicate that some of these highly successful mutations are not inherently the fittest, but represent a random subset of moderately fit mutations that rose to a high frequency due to genetic hitchhiking^42^ or drift.

Overall, our results highlight key evolutionary forces governing the success of mutations in the evolution of SARS-CoV-2, and show that the relative fitness of mutations is largely predictable when considering the intrahost dynamics of mutations and their genetic and phenotypic properties. However, the higher prediction error for the most successful mutations indicates that forecasting the exact set of mutations that will reach a high frequency remains out of reach, for now. Nevertheless, the generalisable patterns inferred by our models enables us to narrow down the list of mutations that are likely to be successful in the evolution of SARS-CoV-2. These could be combined with insights gleaned from other models of mutation or lineage fitness, and from immunological assays, to potentially inform on candidate mutations for future iterations of vaccine targets. Furthermore, we foresee that this approach could be easily ported to other comparable model systems for which large-scale genomic data is available, such as H3N2 Influenza A, which causes seasonal outbreaks globally. There may also be potential to leverage information derived from intrahost genetic variation in very different pathogens, for example to predict the future emergence of antimicrobial resistance in bacteria exposed to drugs.

## Methods

### Data acquisition

We compiled a comprehensive dataset of SARS-CoV-2 genome assemblies. To do so, the metadata, including the list of SAVs, for all SARS-CoV-2 consensus genomes (n=16,798,863) deposited on GISAID^10,11^ was downloaded on 8^th^ July 2024 in the *TSV* file format. We retained only genome entries marked as complete (i.e., >29,000nt) and that were not of low coverage (i.e., >5% Ns), were not isolated from the host genera *Manis* or *Rhinolophus*, were not associated with cell cultures, and had complete collection dates (i.e., year, month and day). For collection months with more than 200,000 genomes deposited, we randomly subsampled to 200,000 genomes, which was adequate to capture the necessary diversity of data while facilitating computational feasibility.

The metadata of all Biosamples hosted on NCBI were downloaded on 21^st^ March 2024 from the FTP site (https://ftp.ncbi.nlm.nih.gov/biosample/) in *XML* file format. This was converted to *TSV* format with a custom Python script that uses the *ElementTree XML API* v1.3.0. All Biosamples whose taxonomy name, Biosample title, isolate or strain fields contained the terms ‘*Severe acute respiratory syndrome coronavirus 2*’, ‘*cov-2*’ or ‘*cov2*’ were retained. Biosamples were linked to their associated sequencing libraries where possible using NCBI’s *entrez-direct* command-line API v21.6. Sequencing libraries were downloaded in the *SRA* file format and converted to *FASTQ* format using the *prefetch* and *fastq-dump* commands, respectively, as part of NCBI’s *sra-tools* API v3.11. For the Early dataset, we considered only Biosamples whose collection year and month were known and between 1^st^ October 2019 and 1^st^ March 2020. For the rest of the datasets, we considered only Biosamples with complete collection dates that corresponded to the following collection months: Alpha (February 2021), Delta (June 2021), BA.1 (December 2021), BA.5 (June 2022), XBB (February 2023), BA.2.86 (December 2023) (Fig. 1b).

We additionally downloaded sequencing data from an independent study designed for investigating the intrahost diversity of SARS-CoV-2 infections^23^ (BioProject accession: PRJEB42623; referred to as the ‘Tonkin-Hill dataset’). These libraries were processed using the same bioinformatic pipeline and filters described below.

### Bioinformatic pipeline for sequencing data

To ensure the homogeneity of sequencing data, we analysed only paired-end Illumina libraries. If Biosamples were associated to multiple sequencing libraries, the sequencing reads were combined prior to further processing. We removed known sequencing adapters, trimmed read ends (<Q20), and removed low quality read pairs (mean base quality of <Q20) using the *BBDuk.sh* utility of the BBMap v39.06 package (<u>sourceforge.net/projects/bbmap/</u>)^43^. We then aligned the quality-filtered read pairs to the CHM13 human reference genome (assembly accession: GCF_009914755.1) using Bowtie2 v2.5.1^44^, retaining read pairs where both members were unmapped. The unmapped read pairs were then re-aligned to the Wuhan-Hu-1 SARS-CoV-2 reference genome (GenBank accession: MN908947.3), retaining only read pairs where both members were mapped. Duplicate reads were removed using the *markdup* utility from Samtools v1.20^45^, to minimise the impact of PCR duplicates on our intrahost diversity analysis. SAV frequencies were estimated from the de-duplicated, high quality reads using the *call codonvar* utility in Quasitools v0.7.0^46^, specifying an sequencing error rate corresponding to Q20 (‘*--error_rate 0.01*’).

Separately, haploid variant calling was performed using the multi-allelic caller (‘*-m*’) in Bcftools v1.20^47^, specifying a minimum mapping quality and base quality of Q30 (‘*--min-MQ 30*’ and ‘*-- min-BQ 30*’, respectively). Consensus genomes were then generated from the variant calls using a custom R script, masking positions where the coverage depth is less than 10, and ignoring indels. Sites that were masked in >10% of genomes were also replaced with Ns. Finally, the re-assembled consensus genomes were assigned PANGO lineages^48^ and variant names using Pangolin v4.3^49^.

### Quality control of sequencing libraries

The diversity of SAVs detected in the sequencing data could be confounded by whether libraries were associated with serial passaging experiments, laboratory contamination, an excess of sequencing artefacts, or erroneous collection dates. In all these cases, we would expect a larger or smaller than expected number of mutations relative to other libraries at each timepoint. For each dataset, we removed sequencing libraries whose consensus genomes had fewer than 1.5 times the lower quartile, or greater than 1.5 times the upper quartile, of the number of SNPs for that dataset. For the Early dataset, we additionally removed libraries that were classified as either Alpha or Beta variant, as these lineages are known to have emerged at later stages in the pandemic so were likely mislabelled (**Supplementary Note 2**). Biosamples in the BA.1 and XBB datasets with no lineage assignments were also excluded as they had a much lower number of SNPs compared to other lineages in circulation within the timeframes of these datasets (**Supplementary Note 2**; **Supplementary** Fig. 2). We additionally removed Biosamples whose re-assembled consensus genomes had >10% Ns or gaps.

### The fitness of mutations

The response variable for all models was the future fitness of a SAV after the sampling timeframe of a particular dataset. We define this as the number of genomes in the GISAID metadata carrying said SAV after the timeframe of the dataset with a one month buffer. Similarly, prior fitness is defined as the number of GISAD genomes carrying said SAV before the dataset timeframe with a one month buffer. For example, the Alpha dataset comprises samples collected in February 2021 and we estimated future and prior fitness as the number of GISAID genomes carrying each SAV between April 2021 and July 2024, and between December 2019 to December 2020, respectively.

We trained and evaluated four cross-dataset models on our datasets (Fig. 2), and two other models involving geographically restricted datasets (**Supplementary Note 2**). Future fitness was estimated using data from specific time windows to maintain independence of the training and testing datasets. Particularly, the cross-dataset experiments involved:

a. Training on Alpha (fitness estimated from after Alpha to before BA.1), testing on BA.1 (estimated from after BA.1 to the end of the entire GISAID timeframe).
b. Training on Alpha (fitness estimated from after Alpha to before Delta), testing on Delta (estimated from after Delta to before BA.1).
c. Training on BA.1 (fitness estimated from after BA.1 to before BA.5), testing on BA.5 (estimated from after BA.5 to before XBB).
d. Training on BA.1 (fitness estimated from after BA.1 to before XBB), testing on XBB (estimated from after XBB to the end of the entire GISAID timeframe).
e. Training on the independent Tonkin-Hill dataset^23^ (fitness estimated using only European GISAID data from June 2020 to before Delta), testing on BA.1 data sampled in the United States (US) (fitness estimated using US GISAID data from after BA.1 to the end of the entire GISAID timeframe).
f. Training on data sampled outside of the US (non-US) Early data (fitness estimated using non-US GISAID data from after Early to before Delta), testing on the US BA.1 data (fitness estimated using US GISAID data from after BA.1 to the end of the entire GISAID timeframe).

### Physiochemical properties and DMS phenotypes

For each SAV, we generated the physiochemical predictors using the *Bio.SeqUtils.ProtParam* module in BioPython v1.84. Changes and absolute changes in physiochemical properties were calculated as *P_alt_* - *P_ref_* and |*P_alt_* - *P_ref_*|, respectively, where *P* represents the physiochemical score of an amino acid. Amino acid charge was calculated at pH 7. Hydropathy was estimated using the Gravy scale^50^, for which more positive scores indicate greater hydrophobicity. BLOSUM62 scores were calculated using the Biostrings v2.70.2 package in R.

Changes in RBD binding (Δlog_10_*K_D,app_*) and expression (Δlog_10_MFI) were obtained directly from the raw data of previous DMS experiments (https://github.com/jbloomlab/SARS-CoV-2-RBD_DMS_variants/blob/main/results/final_variant_scores/final_variant_scores.csv)^4,18^, and represent the log-ratio of the apparent dissociation constant (*K_D,app_*) and mean fluorescence intensity (MFI) of the SARS-CoV-2 RBD with a SAV relative to the wildtype, respectively. Mean antibody escape was also calculated from previous experiments (https://github.com/jbloomlab/SARS-CoV-2-RBD_MAP_Crowe_antibodies/blob/master/results/supp_data/MAP_paper_antibodies_raw_data.csv)^17^. This score represents the escape fraction of the RBD with an SAV relative to the wildtype estimated under global epistasis models^17^, which we averaged across the 10 human monoclonal antibodies assayed. These predictors were assigned a value of −100 for SAVs that did not have any associated DMS estimates, which was the case for all non-RBD mutations. This predictor therefore also provides models with information on whether an SAV is found within the RBD or not.

Relative solvent accessibility (RSA) of the trimeric spike protein structure (PDB:6VXX) was estimated using the Shrake-Rupley method^51^ implemented in BioPython v1.84 and the ‘Wilke’ reference scale^52^. Water molecules were excluded from the analysis. RSA values for the same amino acid residue were averaged across all subunits in the trimer.

### Genetic linkage

Linkage at the intrahost level was calculated as the Pearson’s correlation coefficient, *r*, of the intrahost frequencies of two SAVs across all libraries within a dataset. Pearson’s *r* was only calculated for SAV pairs where there were at least five libraries with non-zero SAV frequencies. Linkage at the interhost level was estimated using the D’ statistic, as follows:

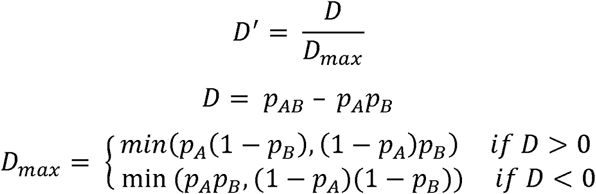

where *p_A_* and *p_B_*, represent the proportion of consensus genomes carrying an SAV at locus A and locus B, respectively, and *p_AB_* represents the proportion carrying both SAVs at locus A and B. For these calculations, we only considered genome entries before the timeframe of each dataset, with a one month buffer (as with the calculation of prior fitness). For example, for the Alpha dataset (February 2021) we consider only genomes collected between December 2019 and December 2020. The *D’* statistic was only calculated for SAVs with a prior fitness of >1000 genomes.

Three types of linkage predictors were generated based on the intrahost and interhost linkage data for each SAV: whether a SAV is strongly linked to at least one other SAV (*D’* or *r*>0.9) is a binary variable; the number of strongly linked SAVs (*D’* or *r*>0.9) is an integer variable ranging from zero to *n* – 1, where *n* is the total number of SAVs considered in the dataset; maximum *D’* or *r* across all pairs involving the SAV is a continuous variable ranging from zero to 1 (negative *D’* or *r* values are set to zero). All linkage predictors that could not be calculated were set to −1 to indicate missingness.

### Model training, hyperparameter optimisation and evaluation

For each dataset, we trained gradient-boosted regressors as part of the XGBoost v2.0.3^26^ module in Python. We optimised the hyperparameters for our models and assessed their test error using a nested 10×10 cross-validation procedure. Briefly, for each iteration of the outer cross-validation loop, a tenth of the data is held out. The remaining nine-tenths of the data is used for the inner cross-validation loop, where an exhaustive search (i.e., grid search) for the best performing combination of hyperparameters is performed. As part of hyperparameter optimisation, 10-fold cross validation is performed to evaluate the performance of models using each combination of hyperparameters, and the best performing model is selected. The held-out tenth of the data in the outer loop is then used to assess the performance of the optimised model, and this process is then repeated for all 10 iterations of the outer loop. We optimised three hyperparameters for our models, particularly the number of trees in the ensemble (n_estimators=100, 300, 500, 700, 900), the maximum depth of each tree (max_depth=1, 2, 3,…, 9), and the proportion of random predictors used to grow each tree in the ensemble (colsample_bytree=0.1, 0.2,…, 1.0). Three main metrics were used for evaluating model performance, the coefficient of determination, *r^2^*, mean absolute error (MAE), and Spearman’s rank correlation coefficient, *ρ*. These metrics are defined by:

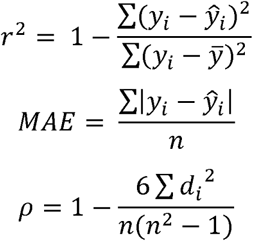

where *y_i_*, *ŷ_i_*, *ȳ*, *n*, and *d_i_*, represent the observed fitness, predicted fitness, mean fitness, number of SAVs considered, and difference in ranks, respectively.

### Model interpretation

To interpret our XGBoost regression models, we used SHAP v0.45.0^27^. Each predictor of a sample is assigned a SHAP value, which corresponds to the change in predicted future fitness given the information of that predictor. SHAP values therefore enable the decomposition of each future fitness prediction into the sum of contributions from each predictor. The relative importance of our predictors were assessed by their mean absolute SHAP values, with higher values indicating more important to the model predictions. The relative importance of groups of predictors (i.e., physiochemical, intrahost diversity, DMS, intrahost linkage and interhost linkage), were assessed by summing the absolute SHAP values for all predictors within a group and dividing this by the number of SAVs considered.

### Statistical analysis and visualisation

All statistical analysis and visualisations were performed using the *stats* and *ggplot2* packages in R v4.3.2. Where applicable, *p*-values where corrected for multiple testing using the Benjamini-Hochberg procedure.

## Acknowledgments

C.C.S.T. is funded by the National Science Scholarship from the Agency for Science, Technology and Research (A*STAR), Singapore. M.E.Z, F.B. and L.v.D. are funded by the European Commission (Horizon 2021-2024, END-VOC Project). L.v.D. is additionally funded by the UKRI Future Leaders Fellowship (MR/X034828/1). Views and opinions expressed are however those of the authors only and do not necessarily reflect those of the European Union or the European Health and Digital Executive Agency. For the purpose of open access, the corresponding author has applied a ‘Creative Commons Attribution’ (CC BY) licence to any Author Accepted Manuscript version arising. The authors acknowledge the use of the UCL Computer Science cluster and associated support services, in the completion of this work.

## Author contributions

C.C.S.T and F.B. conceptualised and designed the study. C.C.S.T. performed all analyses with intellectual inputs from all co-authors. C.C.S.T wrote the manuscript with contributions and edits from all co-authors.

## Data and code availability

All custom code used to perform the analyses reported here are hosted on GitHub (https://github.com/cednotsed/intrahost_prediction). All source data used to perform the analyses here are hosted on Zenodo (https://doi.org/10.5281/zenodo.15255016).

