## Extended Data Figure for "Intrahost dynamics, together with genetic and phenotypic effects predict the success of viral mutations"

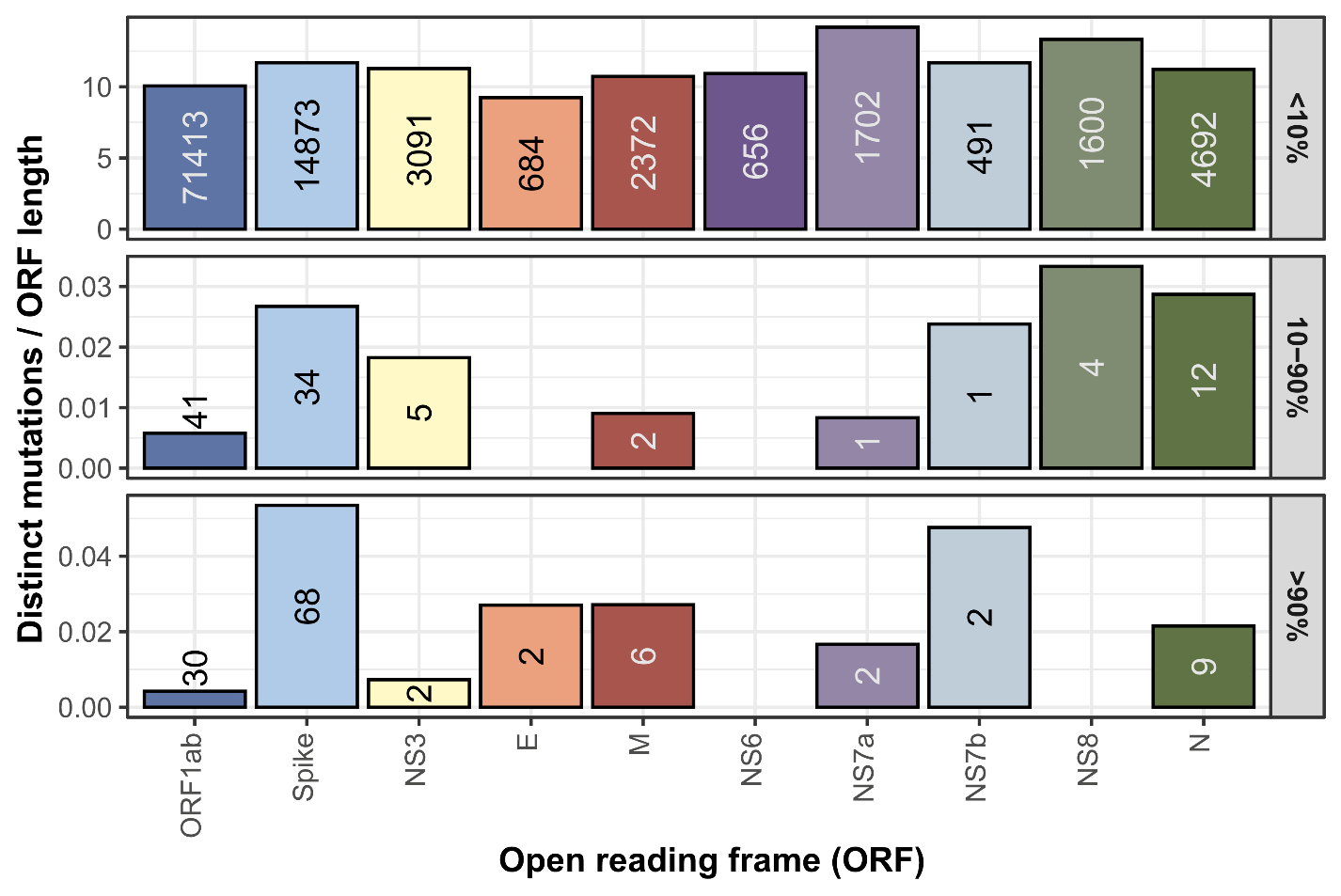

### **Extended Data Figure 1. Distribution of mutations observed in the consensus genomes included in our GISAID dataset.** Number of distinct single amino acid variants (SAVs) observed across the COVID-19 pandemic stratified by open reading frame (ORF) and normalised by ORF length. Plot is generated based on the metadata associated to all complete genomes deposited on the GISAID database from the first submitted sequence up till 8th July 2024.

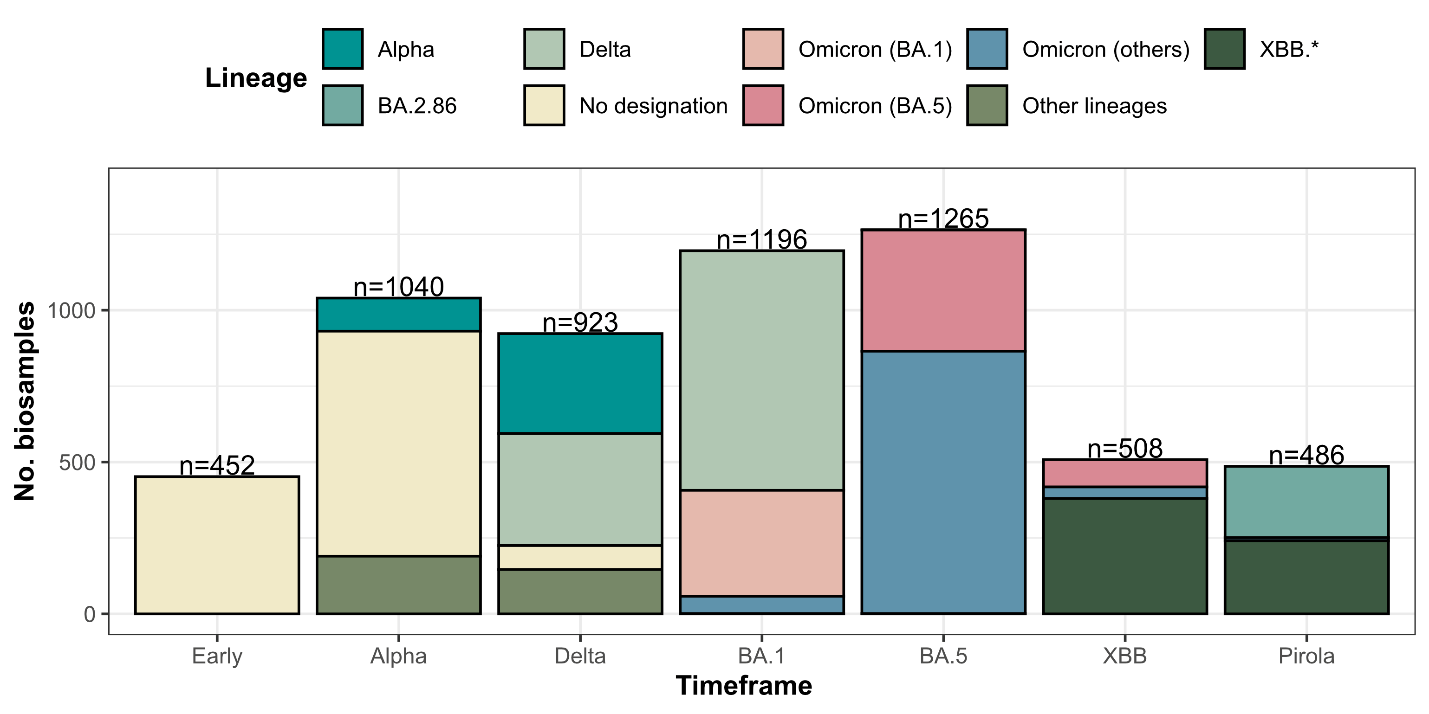

### **Extended Data Figure 2. Sequencing datasets used in this study.** Lineage assignment was performed using PANGO v1.2.6, PUSHER v1.26 and SCORPIO v0.1.12 on consensus genomes that were re-assembled as part of this study.

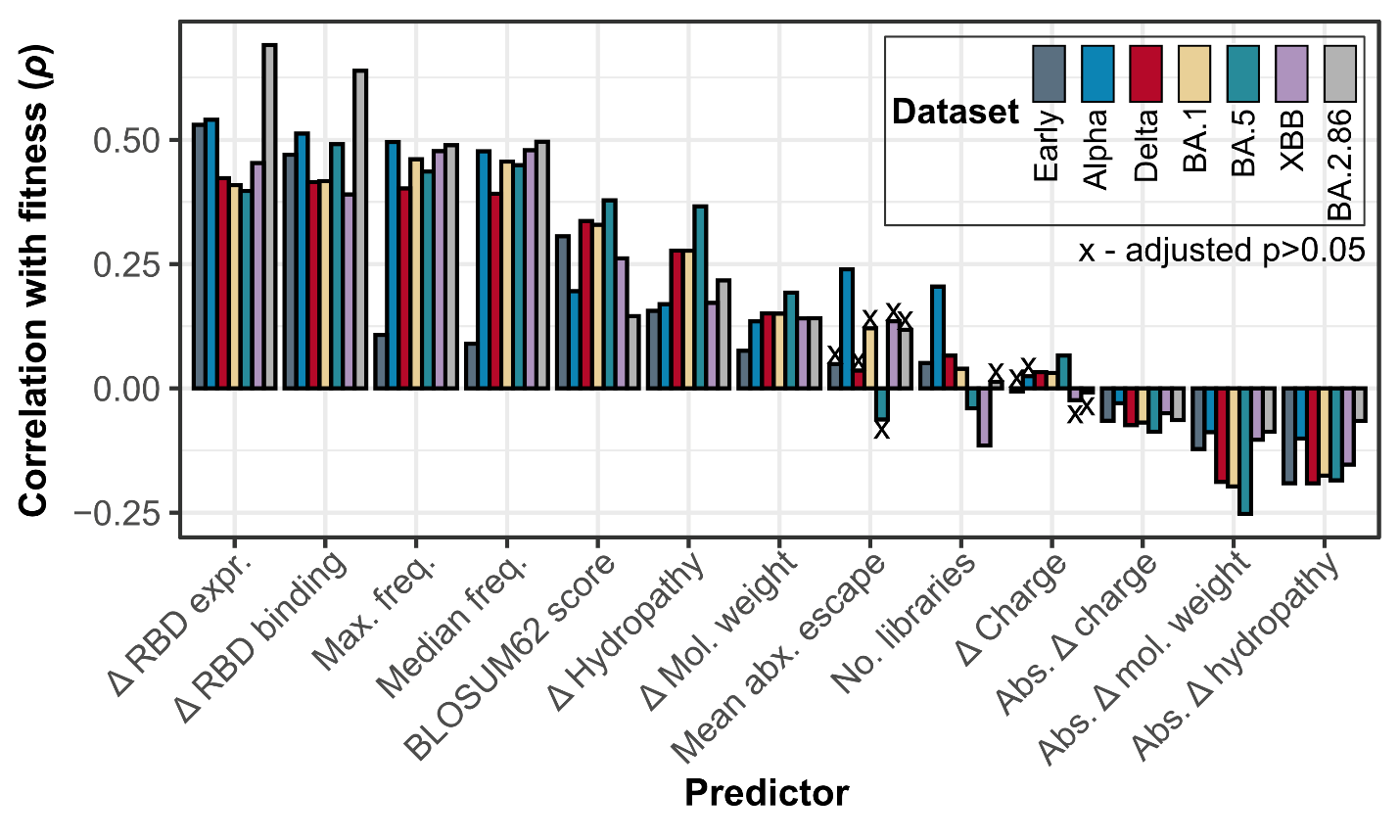

### **Extended Data Figure 3.** Spearman’s correlation between the different predictors included in our modelling approach and the estimated future fitness of SAVs. The statistical significance of each correlation estimate was determined using a two-sided asymptotic t-test and corrected for multiple testing using the Benjamini-Hochberg procedure.

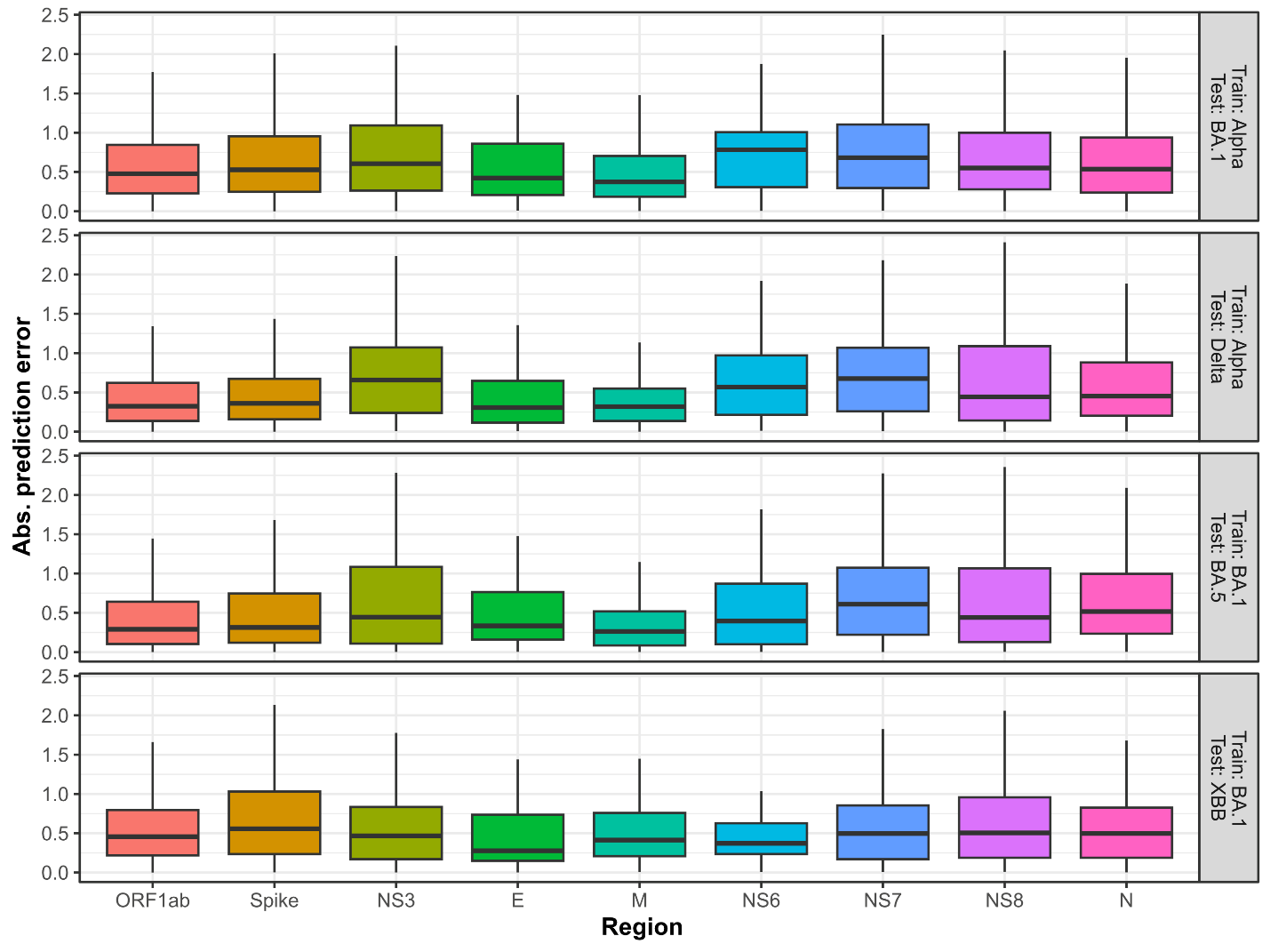

### **Extended Data Figure 4.** Boxplots showing the absolute prediction errors of SAVs in different proteins for our cross-dataset models, which were trained on data from one timeframe and tested on data from another. Boxplot elements are defined as follows: centre line, median; box limits, upper and lower quartiles; whiskers, 1.5x interquartile range. The future fitness values of SAVs were estimated across various timeframes. For example, Alpha-BA.1 refers to the dataset comprising samples collected during the Alpha wave (February 2021) and the fitness values were calculated by considering consensus genomes collected after the timeframe of the Alpha dataset and before the BA.1 dataset (December 2021) with a one-month buffer (i.e., April 2021-October 2021). The timepoint denoted ‘end’ refers to the most recent collection month included in our dataset (i.e., July 2024).

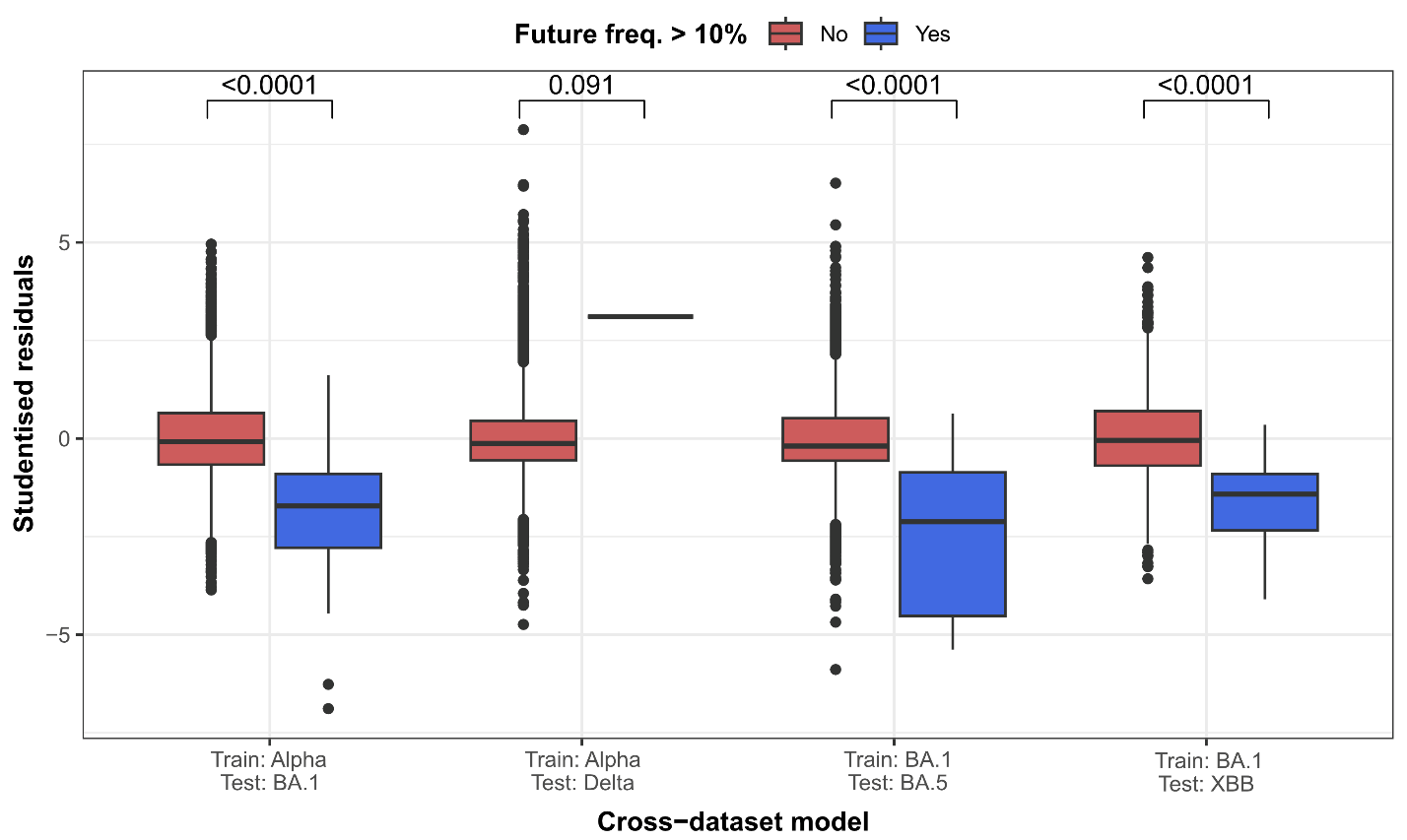

### **Extended Data Figure 5.** Studentised residuals of SAVs, in a linear model of predicted versus observed fitness, for our cross-dataset models. Positive and negative residuals indicate over and under-prediction of fitness values, respectively. Boxplot elements are defined as follows: centre line, median; box limits, upper and lower quartiles; whiskers, 1.5x interquartile range. Differences in distributions were tested using two-sided Mann-Whitney U tests and the corresponding p-values are annotated. The future fitness values of SAVs were estimated across various timeframes as described in **Extended Data. Fig. 4**.

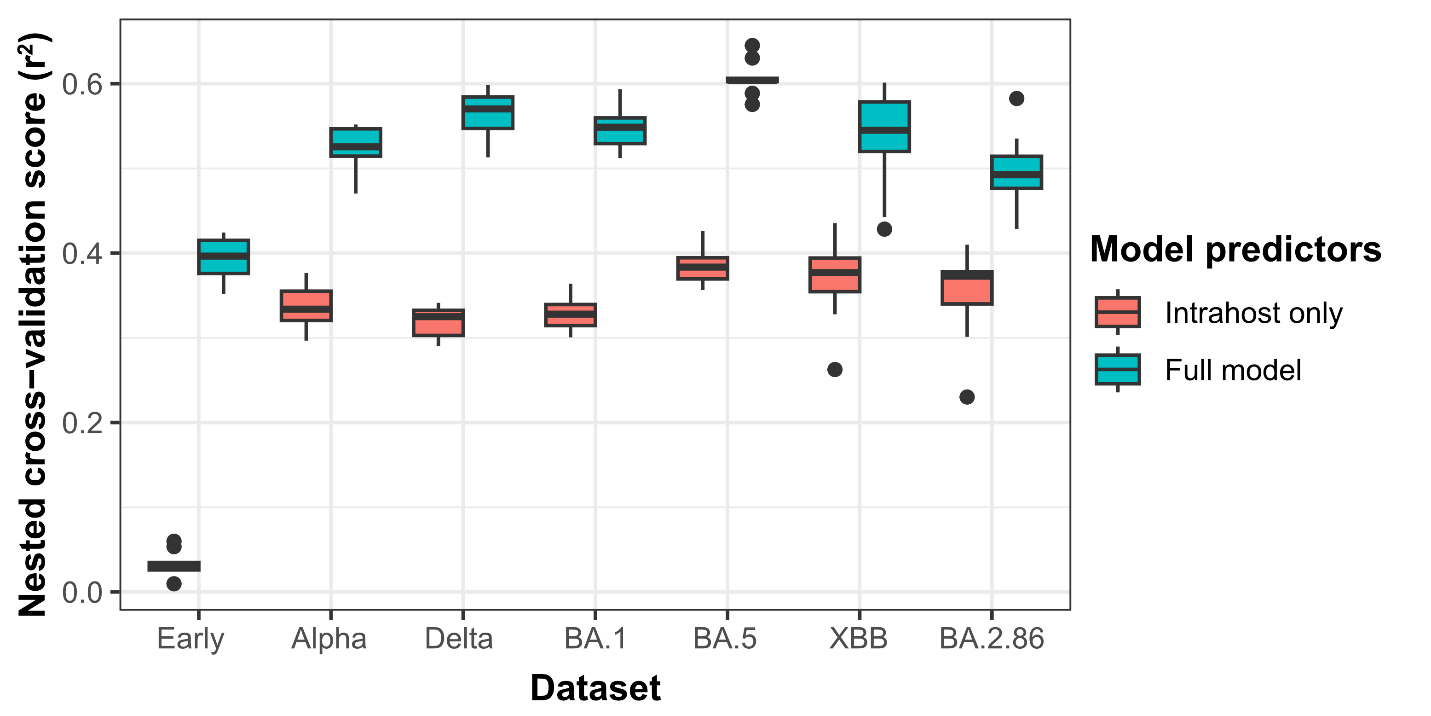

### **Extended Data Figure 6.** Performance of models leveraging only intrahost predictors (orange), and those using all 13 intrahost (teal), physiochemical and phenotypic predictors. Intrahost-only models were trained by randomising all non-intrahost predictor values. Boxplot elements are defined as follows: centre line, median; box limits, upper and lower quartiles; whiskers, 1.5x interquartile range. Differences in the score distributions between model types were tested using two-sided Mann-Whitney U tests. All p-values were highly significant (p<0.0001).

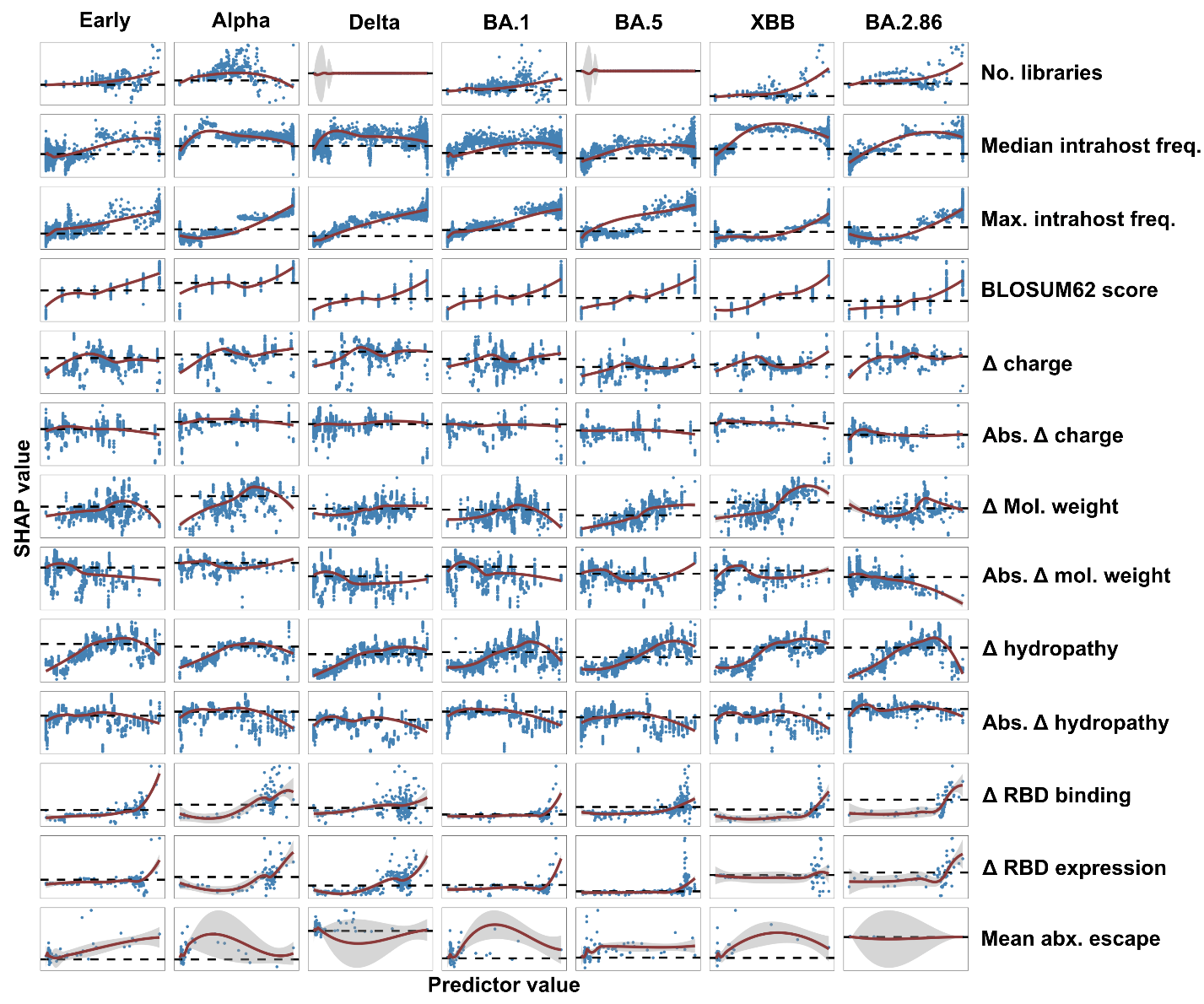

### **Extended Data Figure 7. Relationships between each of the considered predictors and predicted fitness of SAVs.** Scatter plots of predictors and their corresponding SHAP values, showing the predictor-response relationships inferred by the XGBoost regression algorithm. Each point represents the relative contributions of each predictor to the predicted fitness of each SAV (i.e., SHAP value). The red curves and black horizontal dotted lines indicate the LOESS regression smooths and SHAP=0, respectively.

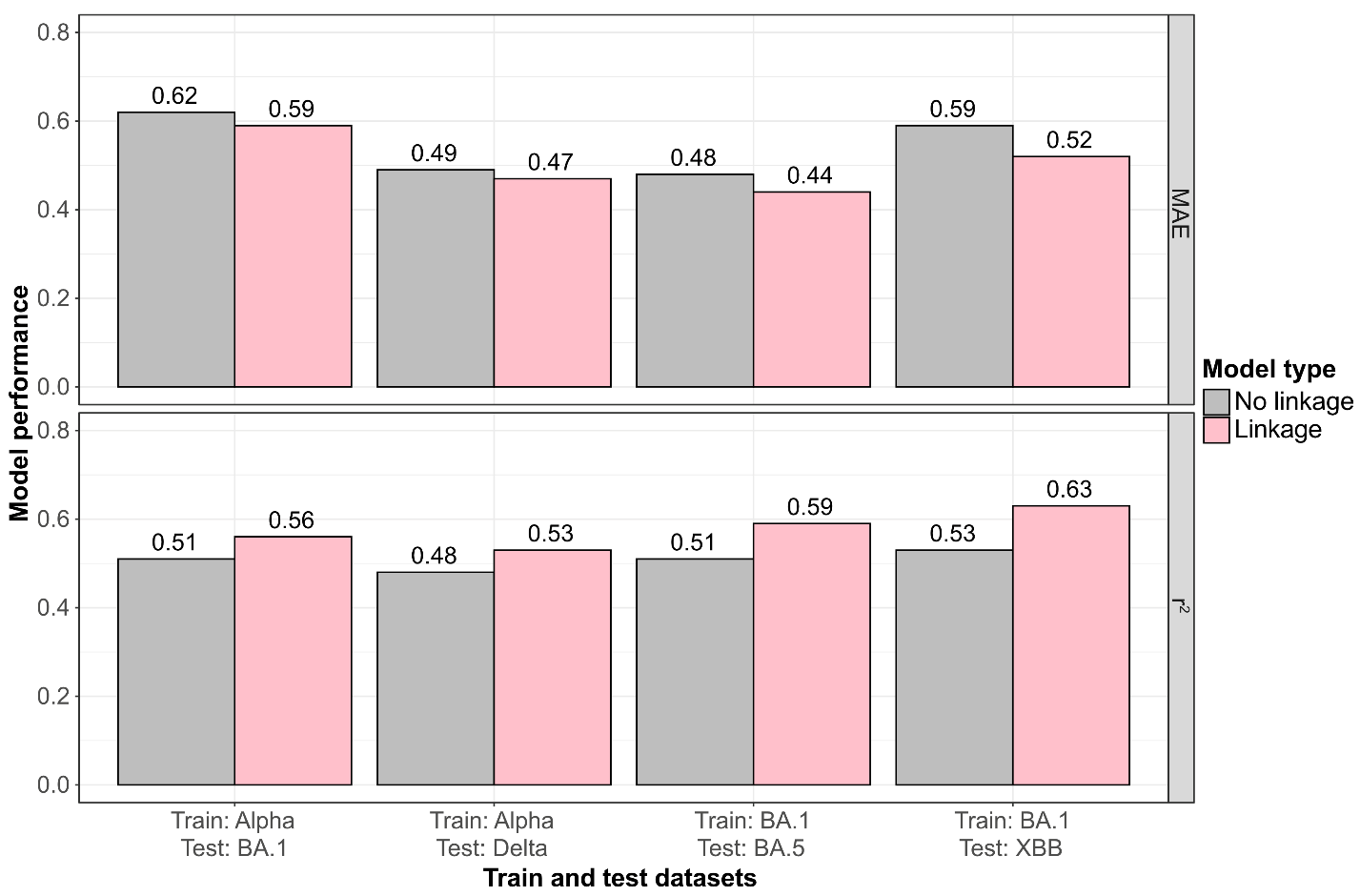

### **Extended Data Figure 8. Model performance when training and testing on different datasets.** Barplots showing the mean absolute error (MAE) and coefficient of determination (*r^2^*) of models trained on various datasets (as given on the x-axis) and tested on other datasets, with or without linkage predictors. The future fitness values of SAVs were estimated across various timeframes as described in **Extended Data. Fig. 4**.
