## Supplementary Note for "Intrahost dynamics, together with genetic and phenotypic effects predict the success of viral mutations"

### **Supplementary Note 1: Mutational fitness estimates are robust to sampling biases**

Genomic databases are inherently biased by sampling and reporting efforts, which may vary greatly by geographical region and over time. Our curated genomic datasets, which include those used to assess intrahost diversity and those used to approximate mutational fitness (i.e., the number of genomes an SAV is found in), are therefore subject to these biases. It is therefore important to assess the effects of sampling biases on our results.

To assess the effects of geographic sampling biases on the mutational fitness estimates, we divided the genome assemblies available in the GISAID dataset into data generated and uploaded from six regions (i.e., Asia, Europe, Africa, North America, South America, Oceania). We then compared the SAV frequencies estimated using data from each region across the 56 months represented by the GISAID dataset. Across all pairwise regional comparisons, we found a consistently high correlation between the SAV frequencies estimated from each regional dataset (minimum Pearsons’s r=0.94; **Supplementary Fig. 1a**). To further investigate whether the impact of (temporal) sampling biases on the fitness estimates changed over the course of the pandemic, we re-performed this analysis but with the GISAID data split at eight-month intervals. We similarly found a high correlation between the SAV frequencies estimated from each regional dataset, but with slightly lower correlations during the first three eight-month intervals (spanning December 2019-November 2021) (**Supplementary Fig. 1b**). These differences may be due to a relatively lower surveillance intensity during the initial years of the pandemic, coupled with regional variation in the implementation and scale of surveillance. However, a model trained on the Early dataset (with future fitness values estimated between May 2020 and April 2021) and testing on the Delta dataset (future fitness estimated from February 2023 to July 2024) showed comparable performance (R^2^=50; MAE=0.62) to models from our other cross-dataset validation experiments (**Fig. 2b**-**d,** see main text). This indicates that the relatively higher sampling bias in the data from the first two years of the pandemic do not considerably impact our model performance.

To further test the effects of sampling biases, we performed a cross-dataset validation experiment where models are trained on data from one region and tested on that from another region. To do this, we constructed an intrahost dataset in the Early timeframe with samples collected outside of the USA to train a model, and tested the model on samples in the Delta timeframe that were collected in the USA. We selected the USA due to the marked intensity of sampling which varied over the course of the pandemic. We additionally re-processed the samples generated and analysed for their intrahost diversity by Tonkin-Hill et al.^1^ (collected between March-April 2020 in the UK)^1^, trained a model using fitness estimates derived from only European consensus sequences, and tested the model on our BA.1 dataset but only considering US samples and consensus genomes. Both models, although trained and assessed on data from different geographical regions, showed comparable performance (Spearman’s ρ=0.70 and 0.68, respectively; both r^2^=0.50) to models from our other cross-dataset validation experiments (**Fig. 2b**-**d**). This indicates that our modelling approach is robust to geographical sampling biases.

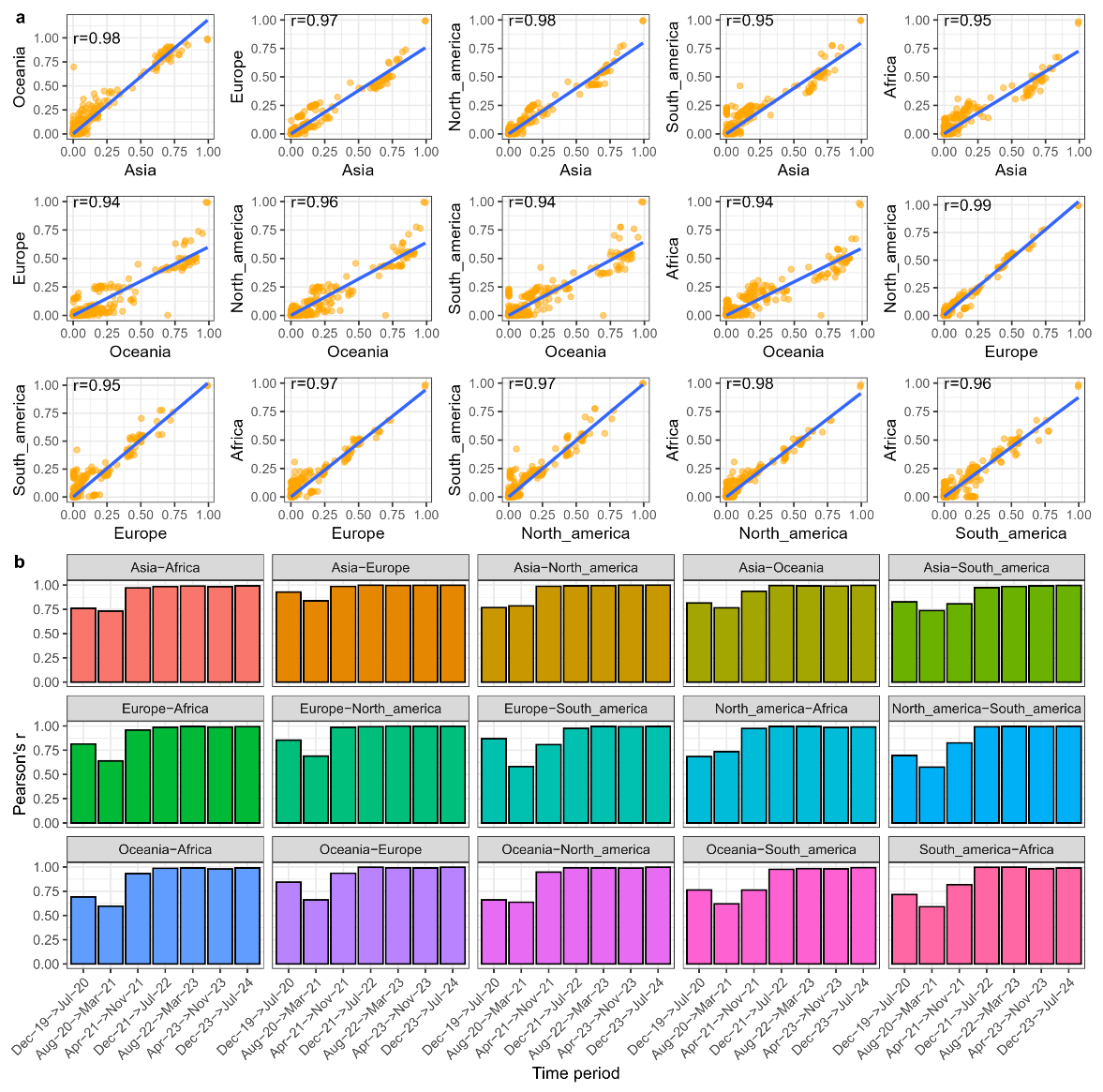

Supplementary Figure 1. *High concordance between regional fitness estimates of mutations. Scatter plots showing all pairwise comparisons of mutational fitness scores estimated using different regional GISAID datasets (a) across all time, and (b) within each eight-month window.*

### **Supplementary Note 2: Quality control of intrahost datasets**

We applied quality control filters to remove libraries potentially associated with serial passaging experiments, laboratory contamination, an excess of sequencing artefacts, or erroneous collection dates. To do so, we reconstructed the consensus genome of all samples and performed lineage classification of these genomes using Pangolin^2^. We then removed any genomes assigned to lineages that were not likely to be circulating during the timeframes of our datasets based on global reporting information (see **Methods**; **Supplementary Fig. 2**). This was done by identifying libraries whose consensus genomes were outliers when assessing the number of single nucleotide polymorphisms relative to the Wuhan-Hu-1 reference (GenBank: MN908947.3) compared to other consensus genomes that were assigned to the same lineage (see **Methods**; **Supplementary Fig. 2**). Additionally, samples that were associated with an outlying number of consensus SNPs were removed (see **Methods**; **Supplementary Fig. 2**). These filters excluded samples that likely had incorrect submission dates. For example, the two Alpha biosamples (SAMEA10053759 and SAMEA10053760) in the Early timeframe both had collection dates of 1^st^ March 2020, but further inspection of their associated consensus genomes on GISAID indicates that these were both collected on 7^th^ February 2021, in line with their Alpha lineage assignment. Additionally, the Beta biosample in the Early timeframe (SAMN18917750) was collected in South Africa on 12^th^ February 2020, but the first reported SARS-CoV-2 infection in Africa is 28 February 2020, so it is most likely that the collection date of this biosample is incorrect.

The median coverage depth, base quality and uniquely mapped reads across the final 5870 samples analysed was, 1260x, Q36, and 312,282 reads, respectively. Further, of these samples, 86% had more than >50,000 uniquely mapped reads. These findings indicate that the sequencing libraries considered were of high quality and had sufficient read depths for reliable estimation of intrahost diversity.

We found that the mean number of subconsensus SAVs per sample across all our quality-controlled samples was 36, which is much higher than those reported by Lythgoe et al.^3^ (mean=0.73), Tonkin-Hill et al.^1^ (mean=9.5), Valesano et al.^4^ (mean=10), Shen et al.^5^ (mean=3.6) and Popa et al.^6^ (mean=11). This is likely due to the fact that our samples are much more geographically diverse and sequenced to much higher read depths compared to those analysed by published studies. Additionally, this difference could also be due to differences in the bioinformatic methods and filters used to identify intrahost SAVs, which was also noted by Lythgoe et al.^3^:

*‘These low levels of SARS-CoV-2 within-host diversity during acute infection are consistent with other reported levels (26, 33) but lower than in some other studies (24, 25), likely reflecting how variants were identified.’*

Importantly, all studies mentioned above called intrahost SAVs using the outputs of per-nucleotide-position SNP callers, which identifies only SAVs arising from single nucleotide changes. Meanwhile, we used a per-read variant caller (i.e., *Quasitools*), which directly identifies SAVs by translating mapped reads, enabling the detection of SAVs arising from more than one nucleotide change. For example, per-position callers will only detect aat🡪aaG but not aat🡪aTG, whereas Quasitools report the frequencies of both. Indeed, when we applied our bioinformatic pipeline to the Tonkin-Hill dataset, the mean number of subconsensus SAVs increased from the reported mean number of 9.5 to 52, confirming Lythgoe and colleagues’^3^ suggestion that the differences in the number of intrahost SAVs identified are likely due to the various detection approaches used.

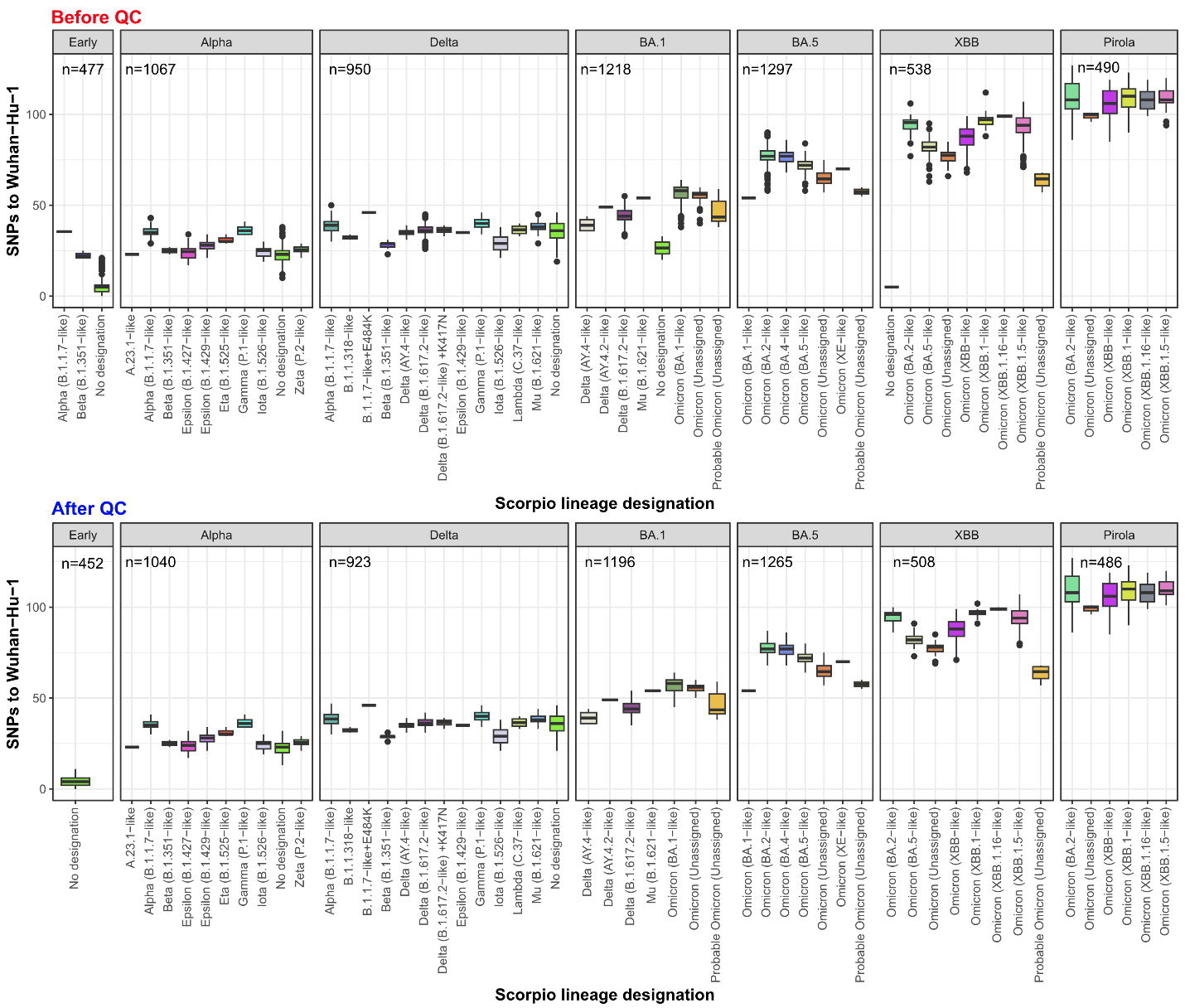

Supplementary Figure 2*. Quality control of sequencing libraries. Distributions of SNPs per PANGO lineage (PANGO v1.2.6) before and after removing libraries that were outliers in terms of the number of SNPs to Wuhan-Hu-1. Alpha and Beta libraries in the ‘Early’ dataset were excluded as these lineages were likely misannotated given they were not observed during the first four months of the COVID-19 pandemic. Libraries with no lineage assignments in the XBB dataset were excluded as they had much lower number of SNPs compared to other lineages in circulation within the timeframe of the dataset.*

### Supplementary Note 3: A**ccounting for cross-contamination of samples**

As noted in previous studies^1,3^, contamination of samples sequenced in the same run may result in the observation of intrahost SAVs that are not endogenous to a sample. We therefore assessed the extent and impact of two major sources of contamination: index hopping and cross-contamination.

*Index hopping*

During multiplex sequencing runs, sequencing reads for a sample may be mislabelled with indices from other samples. The expected index hopping rate is 0.1-2% (<https://emea.illumina.com/techniques/sequencing/ngs-library-prep/multiplexing/index-hopping.html>). On this basis most, if not all, intrahost SAVs arising from index hopping would be filtered out by our intrahost frequency threshold of 3%. As such, we believe the effects of index hopping has been largely accounted for.

*Cross-contamination*

Samples may be contaminated with other samples harbouring different variant lineages during laboratory processing. In the case of severe cross-contamination, some consensus SAVs in the contamination source may appear as consensus SAVs in our samples. The effects of this is likely minimised as a result of our quality control filters (see **Supplementary Note 2**). Additionally, consensus SAVs arising from cross-contamination would also result in phylogenetic conflicts when performing lineage classifications, which is measured as a ‘conflict score’ by Pangolin^2^. In our final quality-controlled dataset, only 212/5870 (4%) of our included samples had conflict scores greater than zero, indicating that severe cross-contamination is not pervasive in our dataset. As phylogenetic conflicts may also arise due to recombination or low diversity early in the pandemic, we opted not to remove these from the final dataset as they may represent genuine genetic diversity.

When cross-contamination is less severe, consensus SAVs in the contaminant source may instead be observed as subconsensus SAVs in samples. To assess the impacts of this, we compared the list of subconsensus SAVs we detected to an existing database of lineage-defining SAVs carried by past variant lineages (including the Variants of Concern; https://github.com/cov-lineages/constellations). If the subconsensus SAVs in our samples were a result of cross-contamination, some samples would carry a large number of subconsensus SAVs associated with the same variant lineage. For example, if a sample were contaminated with a Delta variant sample, we expect to see many Delta-defining SAVs. However, each sample harboured only a small number of lineage-defining subconsensus SAVs (median=0; **Supplementary Fig. 3a**), suggesting these SAVs likely arose de novo. Notably, 1108/5870 (19%) samples harboured multiple lineage-defining subconsensus SAVs (**Supplementary Fig. 3b**). If the SAVs in these samples were a result of cross-contamination, a large proportion would belong to a single contaminant variant lineage. However, the number of lineage-defining subconsensus SAVs found in each sample is strongly correlated (Pearson’s r=0.90) with the number of source variant lineages (**Supplementary Fig. 3b**). This is largely inconsistent with the scenario of cross-contamination from a single variant lineage.

Finally, we tested whether successful SAVs could also be detected in the intrahost diversity of an independent, well-controlled cohort where cross-contamination is minimised. To do so, we applied our sequence processing pipeline and machine learning modelling approach to samples generated by Tonkin-Hill *et al.*^1^ (collected between March-April 2020; henceforth ‘Tonkin-Hill timeframe’). This study was uploaded to BioRXiv on 25 December 2020 (before Alpha reached dominance). As such, we expect minimal cross-contamination from any VoC that emerged after Alpha. The model trained on the Tonkin-Hill timeframe had comparable results to that trained on the Early timeframe (r^2^=0.37 and 0.40, respectively). As with our initial results, we additionally find 59 intrahost SAVs that reach >10% frequency at the consensus level at a median of 19 months after the timeframe of the dataset, the majority of which (53%) were detected in multiple samples (**Supplementary Fig. 4**). These findings further support that our initial observations were not likely due to cross-contamination.

While we cannot completely rule out the effects of contamination, our analyses jointly indicate that our findings are not severely impacted by contamination.

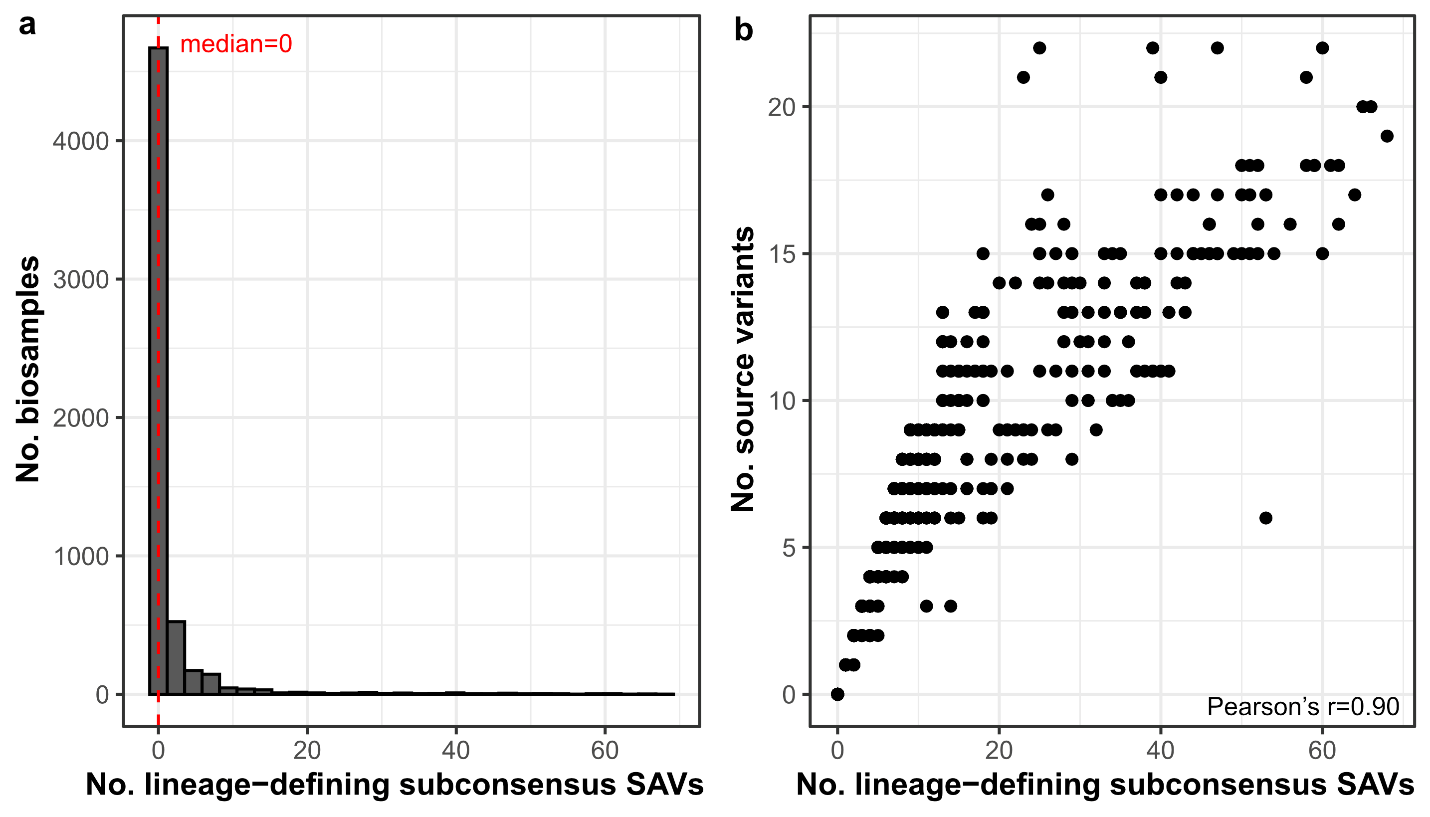

Supplementary Figure 3***.*** *(a) Distribution of the number of source variant lineages for subconsensus SAVs in each sample within our final quality-controlled dataset (n=5885). (b) Number of lineage-defining subconsensus SAVs per sample versus the number of source variant lineages. subconsensus SAVs are defined by an intrahost frequency of <50%. An SAV is defined as lineage defining if it is associated with a variant lineage (including Variants of Concern, Variants Under Monitoring and Variants of Interest), as listed previously (*[*https://github.com/cov-lineages/constellations*](https://github.com/cov-lineages/constellations)*).*

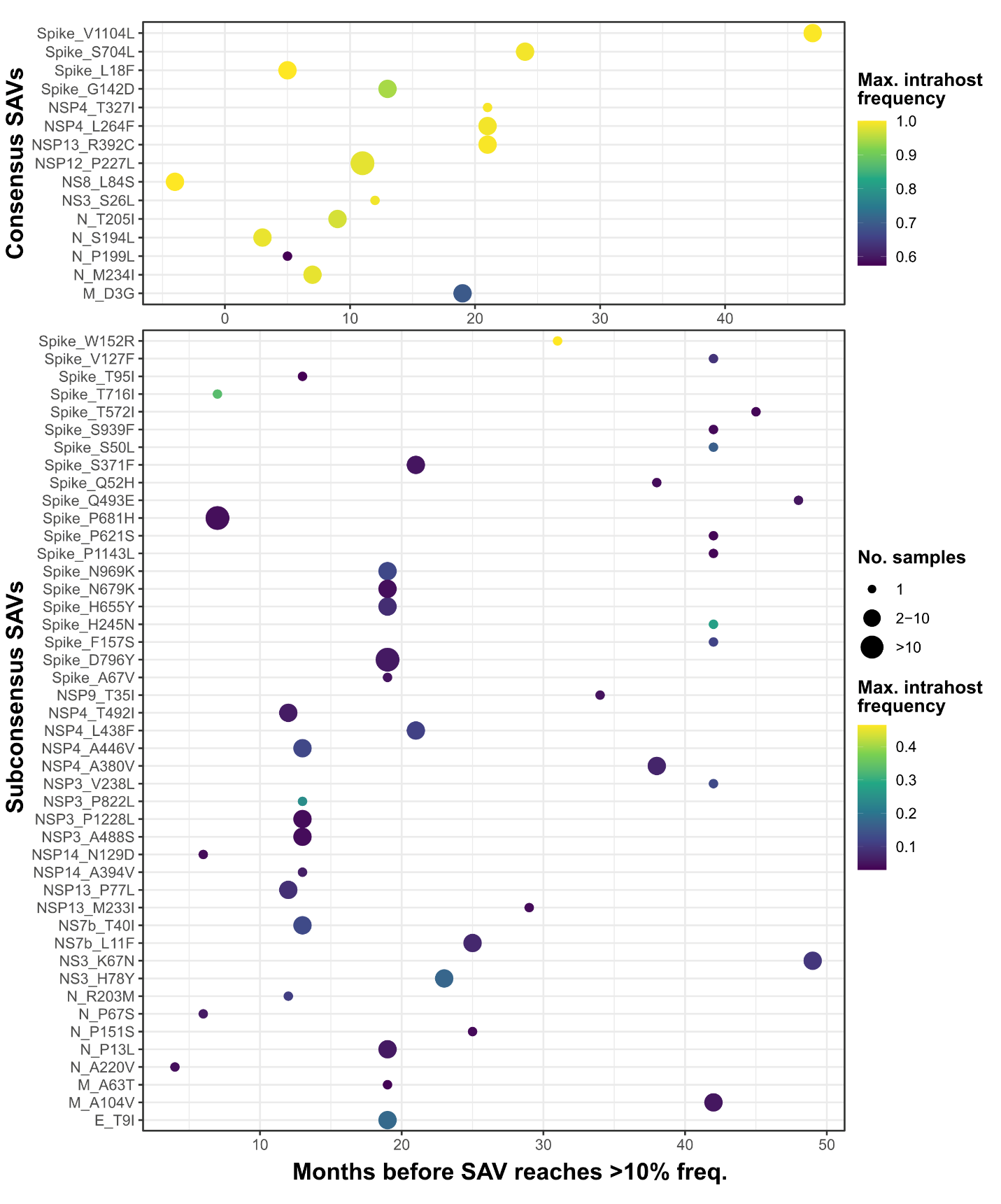

Supplementary Figure 4***.*** *Scatter plots of the intrahost consensus (top) and subconsensus (bottom) SAVs identified in the Tonkin-Hill timeframe, showing the number of months between the end of the timeframe and when each SAV reaches >10% frequency at the consensus level. Size of points indicate the number of samples an SAV is detected in.*

### Supplementary Note 4: Exploring the mechanisms underlying the high fitness of non-conservative SAVs

Both our analysis of GISAID data, and the model interpretation results indicate that large physiochemical changes are generally deleterious. Concordantly, an analysis of the fitness effect scores produced by Bloom and Neher^7^ found that most mutations are either deleterious or neutral (**Supplementary Fig. 5a**), and that more ‘conservative’ SAVs tend to be more fit (**Supplementary Fig. 5b**).

However, we also noted that some non-conservative SAVs can also lead to positive fitness gains. To investigate further, we stratified all SAVs in the GISAID dataset (as in **Fig. 1**) by whether they led to marked physiochemical changes and by whether they reached a high (>10%) population frequency. We found that the high-frequency SAVs that caused large changes in charge and molecular weight are enriched in the spike protein (**Supplementary Fig. 6**). This enrichment of charge-altering SAVs in the spike protein may be linked to adaptation of SARS-CoV-2 and other human coronaviruses for onward transmission in humans. Particularly, previous studies have shown that the spike protein of SARS-CoV-2, HCoV-229E and HCoV-OC43 have evolved to be more positively charged, possibly due to improved binding of the virus to negatively charged carbohydrates in the upper respiratory tract^8,9^. Consistently, we find that the high-frequency SAVs within the upper quartile of charge-change scores confer a much more positive charge change (median=+0.99) than those in the lower three quartiles (median=0) (Mann-Whitney U test, p<0.0001; **Supplementary Fig. 7a**). Additionally, the median relative solvent accessibility of the same high-frequency SAVs is significantly higher for those located at the surface of the spike protein than for those buried within (Mann-Whitney U test, p<0.0001; **Supplementary Fig. 7b**). Overall, these findings hint that perhaps the high fitness of some non-conservative SAVs may be a result of adaptation for onward transmission. However, it is likely that many SAVs are subjected to multiple selective forces – including those linked to functional and structural constraints, and to population immunity – which may be difficult to disentangle.

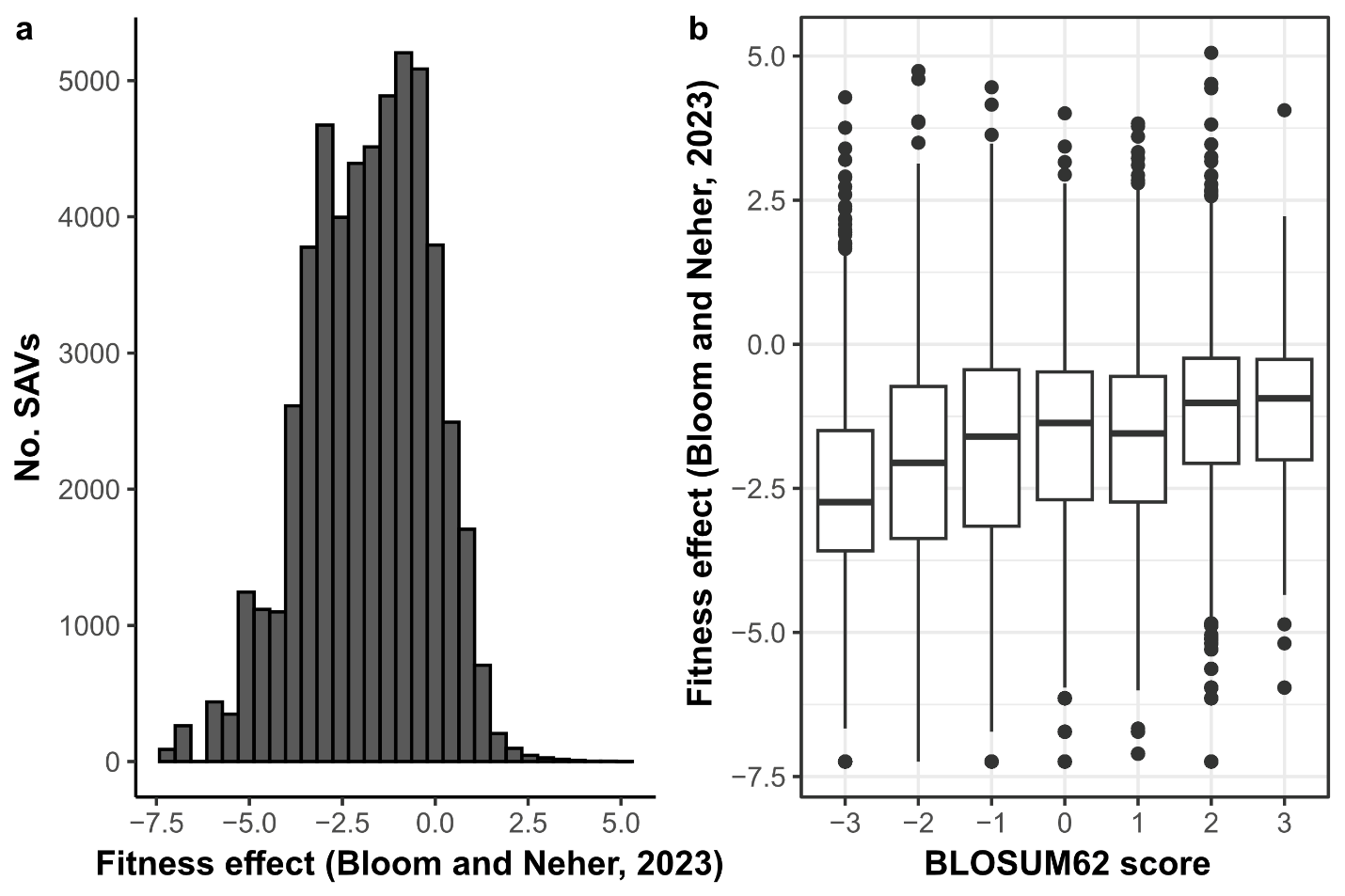

Supplementary Figure 5***.*** *(a) Histogram of fitness effect scores estimated by Bloom and Neher^7^. (b) Positive correlation between mutational ‘conservativeness’ (measured by the BLOSUM62 score) and the fitness effects of SAVs.*

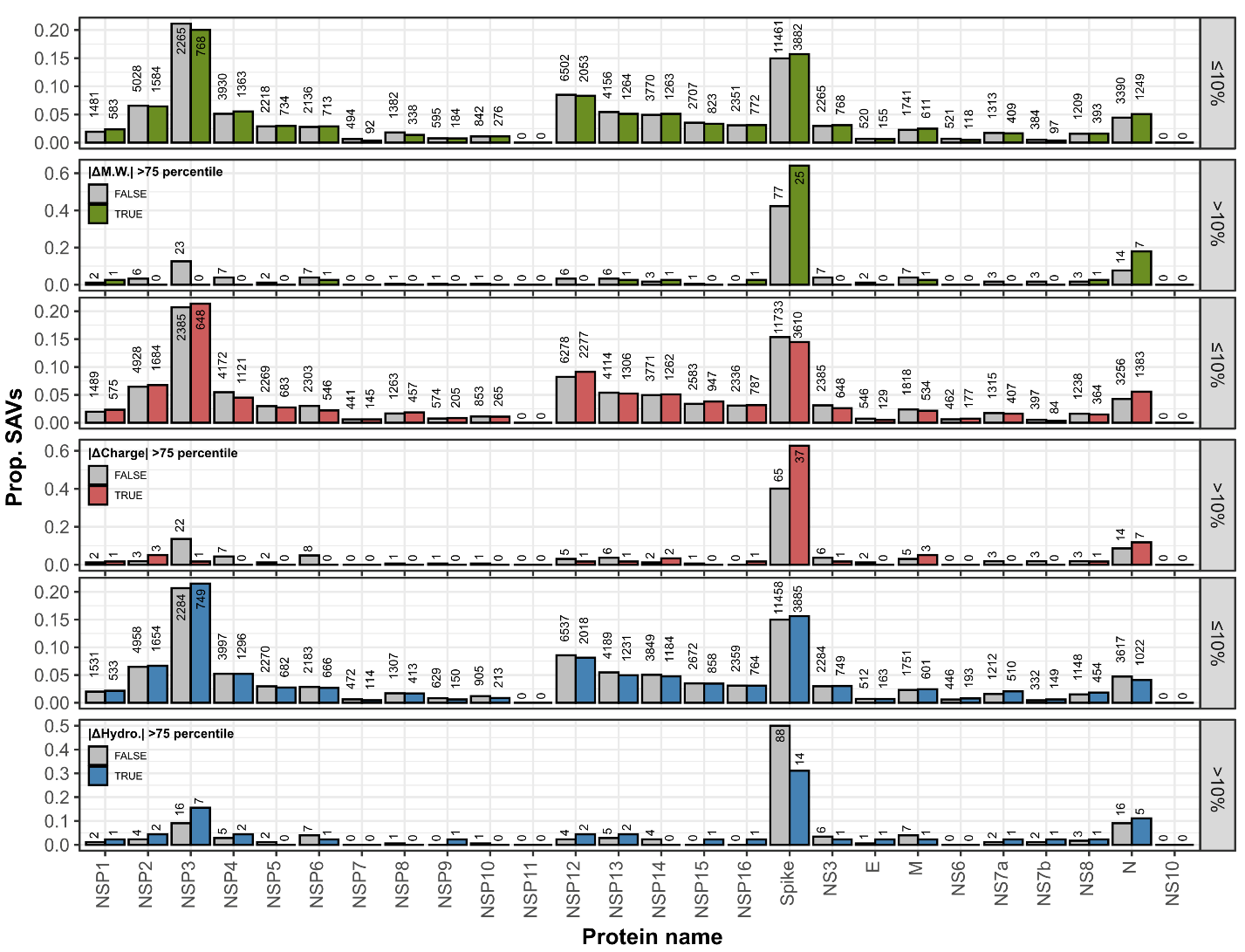

Supplementary Figure 6***.*** *Distribution of the 101,484 SAVs in the GISAID dataset stratified by protein, whether they reached a monthly frequency of >10% at any point, and whether they are on the upper quartile in terms of absolute physiochemical changes (i.e., molecular weight, charge, and hydropathy scores).*

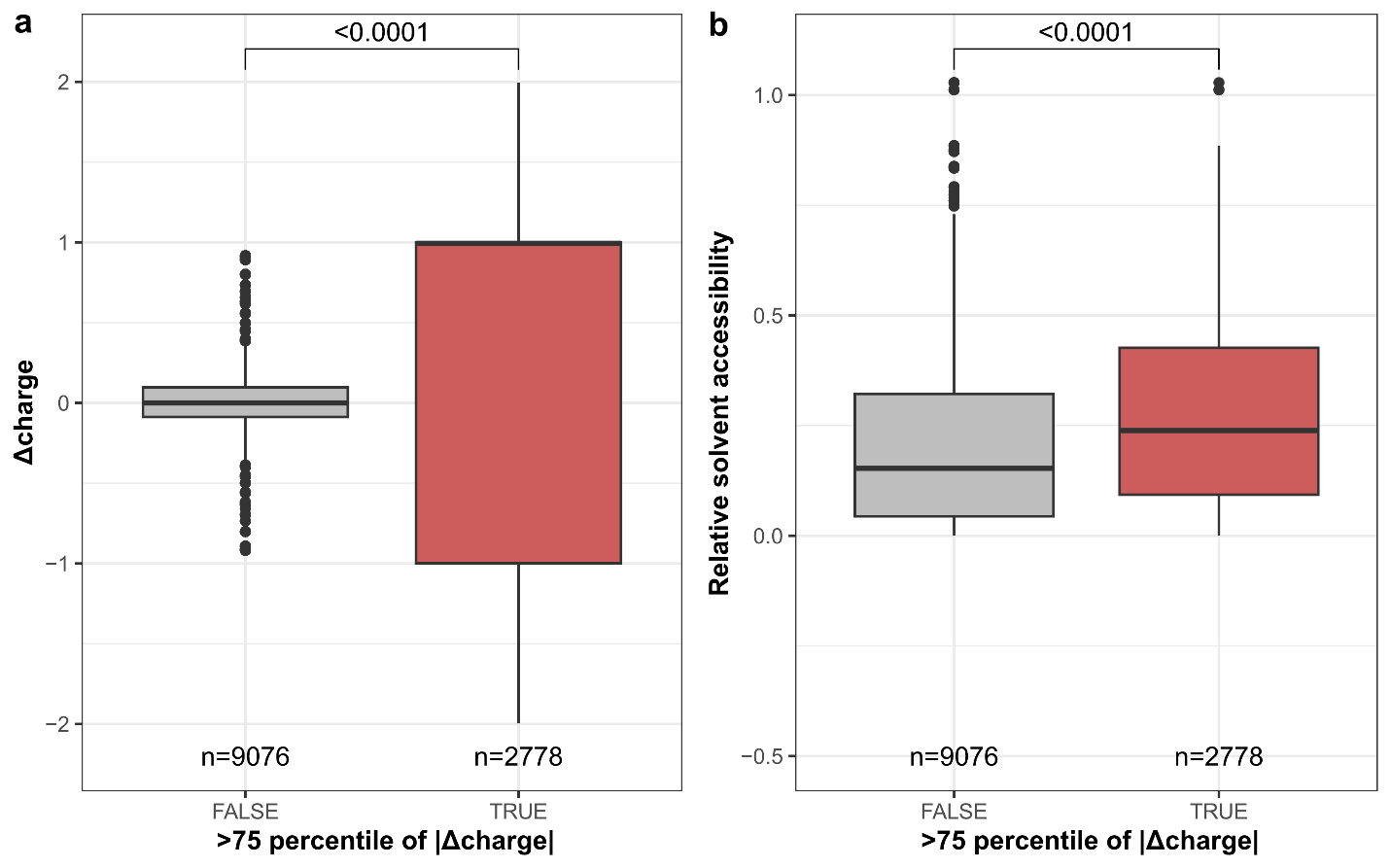

Supplementary Figure 7***.*** *Positively charged SAVs are enriched at the surface of the SARS-CoV-2 spike protein. (a) Estimated charge change and (b) relative solvent accessibility of spike SAVs stratified by whether they fall within the upper quartile of charge change scores. Relative solvent accessibility was calculated from the trimeric, closed-state, uncleaved spike protein structure (PDB:6VXX) using the Shrake-Rupley method implemented in BioPython. Water molecules were excluded. Boxplot elements are defined as follows: centre line, median; box limits, upper and lower quartiles; whiskers, 1.5x interquartile range. Differences in distributions were tested using two-sided Mann-Whitney U tests and the corresponding p-values are annotated.*

### Supplementary Note 5: Selective forces acting on linked SAV pairs

In our results, we noted that interhost linkage is a stronger predictor of mutational success than intrahost linkage. A potential explanation is that the interhost linkage predictors are averaged over a much longer time compared to the intrahost linkage predictors. Additionally, the interhost linkage predictors consider only the SAVs that were successful enough to be transmitted from person-to-person, whereas intrahost predictors consider all SAVs that were generated within the span of a single infection, some of which may be a product of genetic drift and may never be observed at the consensus level. This is supported by the observation that a considerable proportion of strongly linked pairs (r>0.9) comprised only low fitness SAVs (**Fig. 4d**). This is consistent with the hypothesis that the majority of linked SAVs do not confer a significant fitness advantage and are likely a product of genetic drift.

To test this further, we categorised the linked SAVs in our study as under positive, negative or neutral selection based on the HyPhy^10^ selection data generated by Sergei Pond (<https://observablehq.com/@spond/sars-cov-2-global-genomic-selection-2019-aug-2023>). In this resource, codon sites are classified based on the Fixed Effects Likelihood (FEL) method^11^. The proportion of codon pairs classified as diversifying-diversifying and diversifying-neutral was higher for strongly linked codons (D’ or r^2^ > 0.9) across both the intrahost and interhost linkage data (**Supplementary Fig. 8**), indicating that both our intrahost and interhost linkage predictors may capture patterns of co-selection or epistatic interactions. However, the proportion of strongly-linked codon pairs being classified as evolving under neutral selection (i.e., Neutral-Neutral) was 1.8-fold higher in the intrahost data (52%) compared to that in the interhost data (28%) (**Supplementary Fig. 8**). This further indicates that the intrahost linkage predictors likely capture disproportionately more linkage patterns that may result from genetic drift, clouding the relationship between intrahost linkage and mutational fitness. This potentially explains our observation that intrahost linkage predictors have less predictive power than interhost linkage predictors. Nevertheless, it is important to note that the linkage predictors used here – that is, whether the SAV is strongly linked to at least one other SAV (D’ or r>0.9), the number of strongly linked SAVs (D’ or r>0.9), and the maximum D’ or r across all pairs involving the SAV – are imperfect in that they do not consider the specific genetic background the SAV is linked to, the identities of the linked SAVs, or their relative fitness. Designing predictors that would better capture the effects of epistasis in these predictive models is part of our future work.

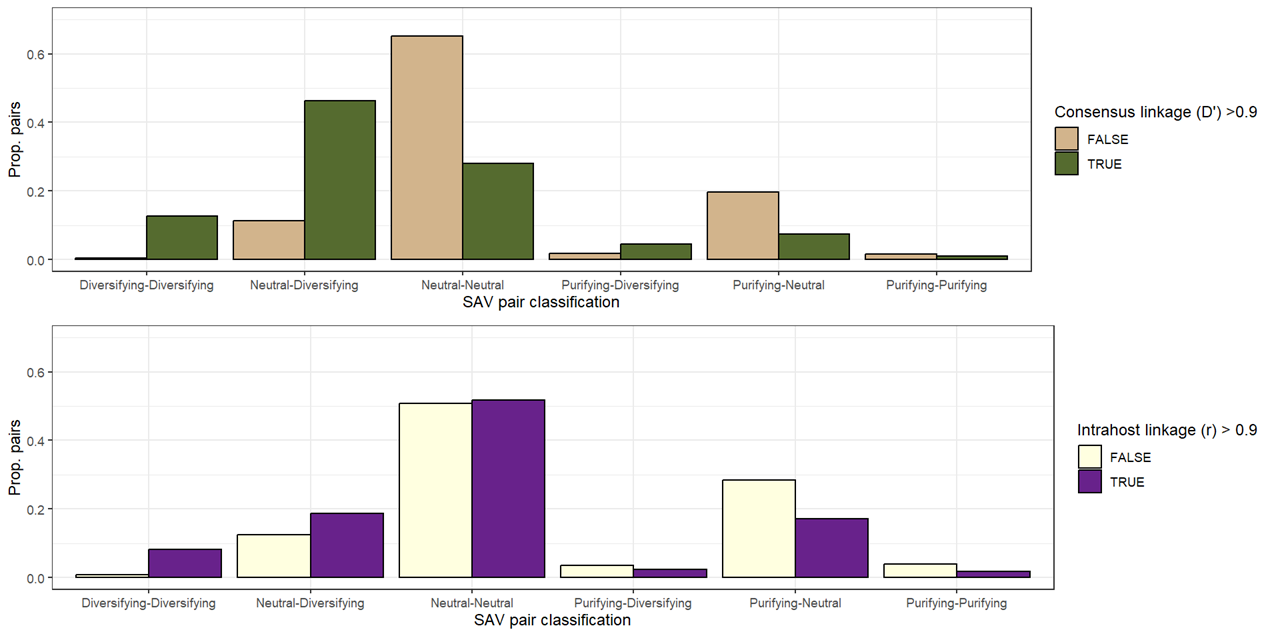

Supplementary Figure 8***.*** *Selective forces acting on SAVs within intrahost and interhost codon pairs.* *Codons were classified as under positive, negative or neutral selection based on selection analysis data produced by Pond 2023 (*[*https://observablehq.com/@spond/sars-cov-2-global-genomic-selection-2019-aug-2023*](https://observablehq.com/@spond/sars-cov-2-global-genomic-selection-2019-aug-2023)*). This data was generated using the Fixed Effects Likelihood (FEL) method*^11^ *in HyPhy*^10^*. The intrahost data shown is based on the consensus D’ estimates calculated between December 2019- December 2023, while the intrahost linkage data is based on the Pearson’s r statistics generated from intrahost SAVs in the BA.1 timeframe.*
